# The complex roles of genomic DNA modifications of bacteriophage T4 in resistance to nuclease-based defense systems of *E. coli*

**DOI:** 10.1101/2022.06.16.496414

**Authors:** Shuangshuang Wang, Erchao Sun, Yuepeng Liu, Baoqi Yin, Xueqi Zhang, Mengling Li, Qi Huang, Chen Tan, Ping Qian, Venigalla B. Rao, Pan Tao

**Author notes:** These authors contributed equally to this work (Shuangshuang Wang, Erchao Sun).

## Abstract

The interplay between defense and counter-defense systems of bacteria and phages is a major driver of evolution of both organisms, leading to their greatest genetic diversity. Bacterial restriction-modification (R-M) and CRISPR-Cas are two well-known defense systems that target phage DNAs through their nuclease activities, whereas phages have developed counter-defense systems through covalent modifications of their genomes. Recent studies have revealed many novel nuclease-containing antiphage systems, which leads to the question of what’s the role of phage genome modifications in countering these systems. Here, we scanned *Escherichia coli* genome sequences available in the NCBI databases and found abundant nuclease-containing defense systems, indicating that phage genomic DNA could be a major target for *E. coli* to restrict infection. From a collection of 816 *E. coli* strains, we cloned and validated 14 systems. Particularly, Gabija and type III Druantia systems have broad antiphage activities. Using wild-type phage T4 and its mutants, T4 (hmC) and T4 (C), which contain glucosyl-5-hydroxymethylcytosines, 5-hydroxymethylcytosines, and unmodified cytosines in the genomic DNA respectively, we revealed the complex roles of genomic modification of phage T4 in countering the nuclease-containing defense systems other than simply blocking the degradation of genomic DNA by nuclease.

## INTRODUCTION

The genomic DNAs of phages, particularly *Escherichia coli* and *Salmonella enterica* phages, are hypermodified, and more than 21 modified nucleotides were identified so far [1, 2]. Many phages encode their own enzymes participating in the genome modifications even though, typically, they have small genomes [3, 4]. This indicates the important roles of such modifications in the life cycle of phages, which has not been fully understood yet. Previous studies have demonstrated genomic DNA modification is the main mechanism of phages to resist restriction-modification (R-M) defense systems of bacteria, which cleave phage DNAs at specific sites while protecting their own genomes [5]. Indeed, phages and their hosts are locked in the endless battle, which drives the evolution of both sides. Phages depend on the host bacteria for reproduction, which usually leads to lysis of infected cells, whereas bacteria must evolve strategies such as the R-M systems to defend themselves against slaughters. Phages in turn have evolved counter-defenses such as the modification of genomic DNA to resist the nucleases of the R-M systems.

One well-studied example is phage T4, of which the genomic DNA contains glucosyl-5-hydroxymethylcytosine (ghmC) instead of cytosine [6, 7]. During phage T4 DNA replication, the 5-hydroxymethyl dCTP, instead of dCTP, is incorporated into the newly synthesized DNA. The 5-hydroxymethylcytosines (hmC) are further glucosylated by phage-encoded α- and β-glucosyltransferase, making wild-type T4 genomic DNA resistant to almost all types I-III R-M systems [8-11]. However, the remaining type IV R-M systems specifically target T4 phages with different genome modifications, for instance, the GmrSD system cleaves ghmC-but not hmC-modified T4 genomic DNA [12] whereas the McrBC system specifically targets phage T4 mutant containing hmC DNA [13].

Recently, we and others found that the ghmC modification also confers T4 phage variable resistance to different types of CRISPR-Cas systems [14-17], the acquired immune systems of bacteria against phages through assembly of Cas-crRNA complexes that specifically recognize and cleave phage genomic DNA [3, 18]. However, bacteria might express different Cas-crRNA complexes targeting independent sites on phage genome to more effectively defend themselves against the phage with modified genome [14, 19]. Additionally, the partial resistance of phage T4 to CRISPR-Cas systems not only allows survival of some *E. coli* cells but also accelerates the evolution of the phages [20-22], highlighting the complex roles of genomic DNA modifications in the coevolution of phages and their hosts.

Other than R-M and CRISPR-Cas, recent studies have revealed many novel antiphage systems that contain nuclease domains such as Zorya, Septu, Gabija, Shedu, Druantia, SspABCD-SspE, STAND, qatABCD, mzaABCDE, to name a few [23-26]. However, their defense mechanisms against phages are not clear. Although all contain at least one nuclease domain, it is unknown for most of the systems whether this domain is necessary for their antiphage activities. Given the fact that *E. coli* phage DNAs are hypermodified [1, 2], it is interesting to know whether all these defense systems exist in *E. coli* and what’s the roles of phage genome modifications in countering these systems.

In this study, we systematically analyzed all the *E. coli* genome sequences available in the NCBI databases and identified 6 new *E. coli* defense systems, which share similar domain structures with recently identified antiphage systems containing nuclease domain in other bacteria. We cloned all these systems, as well as recently identified 8 *E. coli* antiphage systems containing nuclease domains, from a collection of 816 *E. coli* strains maintained in our laboratory. Divergent defense effects of these systems were observed against both wild-type phage T4 and two mutants, T4 (hmC) and T4 (C), which contain hmC-modified and unmodified cytosines respectively in their genome. Our results revealed the complex roles of genomic modification of phage T4 in countering the nuclease-containing defense systems other than simply blocking the degradation of genomic DNA by nuclease.

## RESULTS

### Identification of antiphage systems with nuclease domains in *E. coli*

There are 20 types of nuclease-containing defense systems that have been identified as of February 2021 from a variety of bacteria and archaea, including 12 systems identified in *E. coli* (Table S1). To determine whether the remaining 8 defense systems exist in *E. coli*, we downloaded all the 2,289 *E. coli* genomes available in NCBI till October 2020. The open reading frames of each genome were annotated and analyzed with HMMER [27]. We identified 6 potential defense systems that share similar domain structures with Gabija, Septu, Shedu, mzaABCDE, type III Druantia, and Ago respectively (Fig.1A). However, we didn’t find DISARM and type III AVAST systems in the 2,289 *E. coli* genomes. The *E. coli* defenses systems Gabija, Septu, Shedu, and type III Druantia showed very low homology to the corresponding systems reported in other bacteria, wherea *E. coli* mzaABCDE shared more than 90% sequence similarity with *Salmonella enterica* (Table S2)

**Fig. 1.**
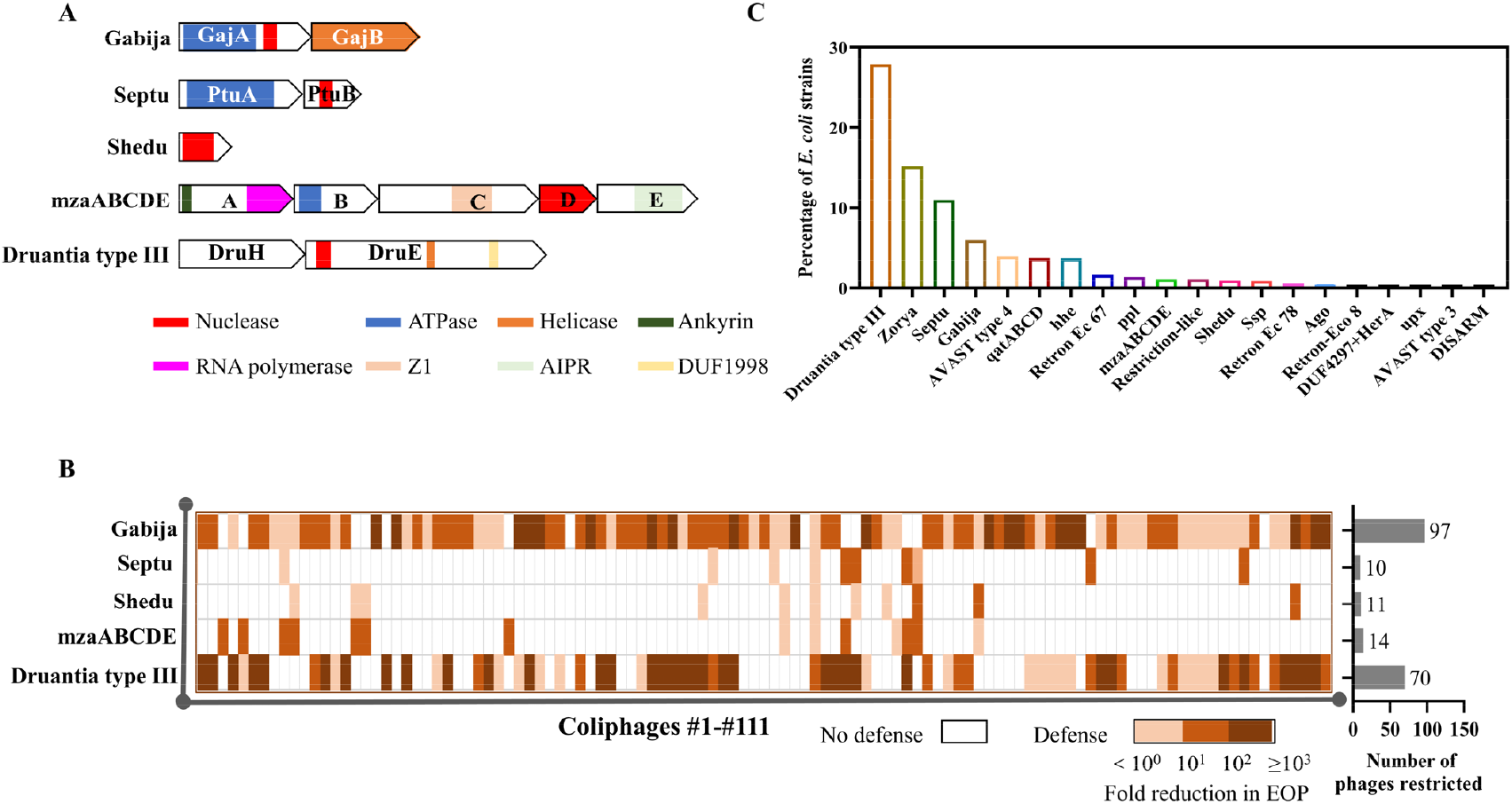
Identification of antiphage systems with nuclease domains in *E. coli*. **(A, B)** Identification and validation of *E. coli* Gabija, Septu, Shedu, mzaABCDE, and type III Druantia systems. Domain organization **(A)** of each system is shown, and their defense activities **(B)** were determined using 111 different phages as described in Materials and Methods. **(C)** The abundance of all 20 nuclease-containing defense systems in the 2,289 *E. coli* genomes.

Using a collection of 816 *E. coli* strains maintained in our laboratory, we were able to clone 5 of the 6 potential defense systems (Fig.1A and Table S3). The defense activity of these systems was determined using *E. coli* MG1655 against 111 different phages. As shown in Fig.1B, Gabija and type III Druantia have broad antiphage activities with more than 10^3^-fold protection against cell lysis for many phages, and the remaining systems also have defense activities although with narrow-spectrum.

Analysis of the abundance of all 20 confirmed defense systems containing nuclease domain in the 2,289 *E. coli* genomes showed that type III Druantia is the most abundant system followed by Zorya, Septu, Gabija, AVAST type 4, qatABCD, and hhe, whereas upx, DUF4297+HerA, Ago, Retron-Eco8, and AVAST type 3 are rare (Fig. 1C). More than half of the strains (1267) strains contain at least one nuclease-containing system, and 10 strains even contain 2 copies of the same system (Table S4), and 424 strains contain more than one nuclease-containing system (Fig. S1). These results demonstrated that divergent nuclease-containing defense systems are employed by *E. coli*, in addition to the well-known R-M systems. The abundance of such systems indicated that phage genomic DNAs could be the major targets for *E. coli* to restrict infection, therefore, it is not surprising that phage genomes are hypermodified.

To determine the roles of phage genome modifications in countering nuclease-containing systems, we, therefore, screened our *E. coli* collections and were able to clone 14 systems (Table S5). Their defense activities were determined with wild-type phage T4, T4 (hmC), and T4 (C) respectively, and discussed in the following sections.

### Genome modifications of phage T4 cannot abolish antiphage activities of Septu, SspBCDE, and mzaABCDE

To determine the anti-defense effects of genomic DNA modifications against the nuclease-based defense systems, we firstly deleted *Alfa-gt* and *Beta-gt* genes encoding α- and β-glucosyltransferase respectively to generate a T4 mutant, T4 (hmC), which contains the hmCs within its genomic DNA (Fig. S2A and Table S6). The mutant T4 (C), which contains unmodified cytosines, was constructed by deleting gene *56* (Fig. S2B) [28]. The modifications of genomic DNAs of wild-type phage T4 and two mutants were confirmed by incubating phage genomic DNAs with divergent nucleases, which degrade DNAs with different modifications as reported previously (Fig. S2C) [12, 15].

Plaque assays revealed that Septu, mzaABCDE, and SspBCDE systems confer *E. coli* cells various degrees of protections against wild-type T4 and two mutants, T4 (hmC) and T4 (C) (Fig. 2A, B). Both Septu and mzaABCDE systems showed similar antiphage activities against T4 (hmC) and T4 (C) phages, with 10^3^- to 10^4^-fold protection against cell lysis. However, defense activity of mzaABCDE system decreased more strongly compared with Septu when wild-type phage T4 (ghmC) was used for infection (10^4^-fold versus 10-fold reduction) (Fig. 2B). SspABCD–SspE system was identified recently in *Vibrio cyclitrophicus, E. coli*, and *Streptomyces yokosukanensis*, which restricts phages through introducing nicks to the genomic DNAs by nickase SspE [24]. Proteins SspA, B, C, and D work together to replace oxygen atoms of genomic DNA backbone with sulfurs in a sequence-specific manner, known as phosphorothioate (PT) modification, which stimulates the nickase activity of SspE. Both PT modification and SspE of *V. cyclitrophicus* are necessary for its antiphage activity [24, 29]. Interestingly, we found *E. coli* does not have *SspA* gene (Fig. 2A), and SspBCDE cassette cloned from our *E. coli* collection is active against wild-type T4 and the mutants (Fig. 2B). The antiphage efficiency depends on the modification degrees of phage genomic DNA, with 10^5^-fold, 10^4^-fold, and 10^3^-fold reductions for phages T4 (C), T4 (hmC), and T4 (ghmC), respectively (Fig. 2B). This revealed that PT modification is not necessary for *E. coli* SspBCDE system to restrict T4 infection, indicating the different defense mechanism of *E. coli* SspBCDE from *V. cyclitrophicus* SspABCD–SspE. Taken together, these results showed the sensitivities of defense systems Septu, mzaABCDE, and SspBCDE on different genome modifications were quite different even though they can protect *E. coli* cells against all three phages, indicating the mechanism differences among these systems.

**Fig. 2.**
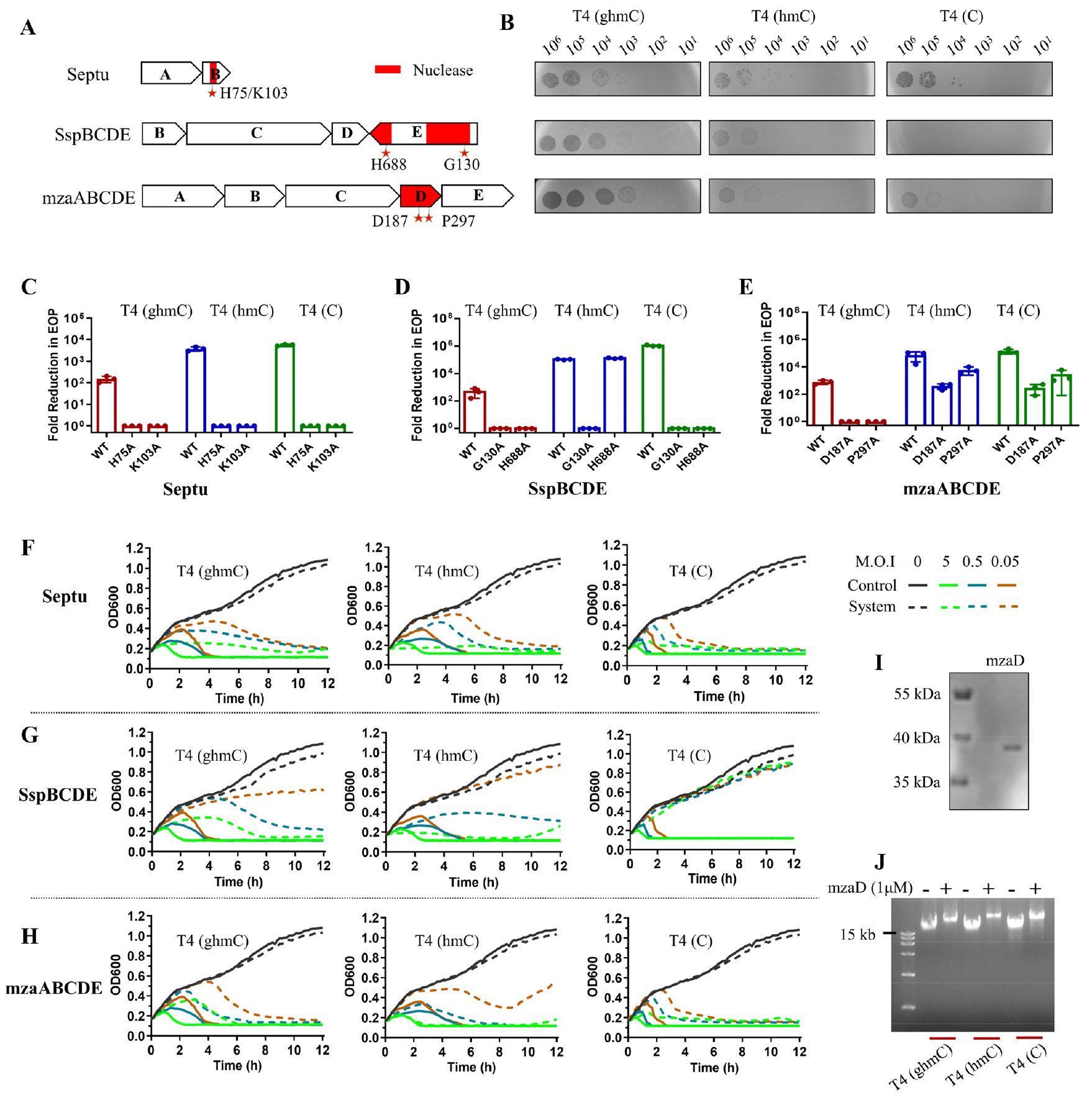
Genome modifications of phage T4 cannot abolish antiphage activities of Septu, SspBCDE, and mzaABCDE. **(A)** Genetic compositions of *E. coli* Septu, SspBCDE, and mzaABCDE systems. The nuclease domains of each system were highlighted with red, and the putative active sites or the conserved amino acids in nuclease domain were indicated with red starts. **(B)** The representative results of plaque assays showing the defense activities of each system against T4 (ghmC), T4 (hmC), and T4 (C). The amount of phages (10-10^6^ pfu) used for plaque assays were indicated on top of each panel. The antiphage activities of wild-type and mutated Septu **(C)**, SspBCDE **(D)** and mzaABCDE **(E)** systems against phages T4 (ghmC), T4 (hmC), and T4 (C) were presented as fold reduction in efficiency of plating (EOP) (see Materials and Methods for the details). Data were represented as mean ± S.D. of three independent assays. **(F-H)** Growth curves of *E. coli* DH10B cells expressing Septu **(F)**, SspBCDE **(G)**, and mzaABCDE **(H)** after infection with T4 phages at different multiplicity of infection (MOI=0, 0.05, 0.5, and 5). *E. coli* DH10B cells without these defense systems were used as controls (solid lines). The nuclease domain-containing protein, mzaD, of mzaABCDE was purified **(I)**, and its nuclease activity on genomic DNAs of phages T4 (ghmC), T4 (hmC), and T4 (C) was determined **(J)**.

To determine whether the nuclease activity is necessary for the defense, we mutated the putative active sites or the conserved amino acids in nuclease domain of each system to alanine (Fig. 2A and C-E). Mutations at sites H75A and K103A completely abolished defense activity of Septu against phage T4 regardless of genomic DNA modifications (Fig. 2C). Interestingly, a single mutation H688A in the conserved EHxxP motif of SspE nickase catalytic center completely abolished the defense activities of SspBCDE against T4 (C) and T4 (ghmC), but did not affect the activity against T4 (hmC) (Fig. 2D). Mutation G130A in the DGQQR motif of NTPase catalytic center of SspE, which was shown to abolish its PT-sensing GTPase function but enhance nicking activity [24], abolished the defense activity of SspBCDE against wild-type T4 and two mutants. These results indicated that SspBCDE might employ different mechanisms to inhibit phage T4 and mutants, and nickase activity of SspE is not necessary for anti-T4 (hmC) activity. Similarly, mutations at sites D187A and P297A only abolished the antiphage activity of mzaABCDE system against T4 (ghmC), but not against T4 (C) and T4 (hmC) (Fig. 2E). These results indicated that both SspBCDE and mzaABCDE systems might employ different mechanisms to restrict T4 phages with different modifications, highlighting the complex roles of phage genome modifications in counteracting bacterial defense systems.

We, therefore, determined the growth kinetics of defense system-containing *E. coli* cells infected by different phages at different multiplicity of infection (MOI) (Fig. 2F-H). The growth kinetics at an MOI of 5 showed that *E. coli* cells expressing SspBCDE system had a lower optical density at 600 nm (OD600) at the first 90 min after infection with T4 (hmC) compared with control *E. coli* cells, indicating the SspBCDE-mediated growth arrest (Fig. 2G, middle panel). Interestingly, it happened only when *E. coli* cells were infected with T4 (hmC) but not T4 (ghmC) and T4 (C), indicating the specific mechanism of SspBCDE against T4 (hmC), which was also observed in our mutation experiments (Fig. 2D). Similar results were also observed for the Septu system (Fig. 2F, middle panel). Growth kinetics of *E. coli* cells expressing mzaABCDE did not show evidences of growth arrest, no matter which T4 phage was used for infection at varying MOI (Fig. 2H). To determine whether mzaABCDE directly targets T4 genome DNAs through its nuclease activity, we expressed mzaD protein (Fig. 3I). Interestingly, we found mzaD can cut genomic DNAs of T4 (C), T4 (hmC), and T4(ghmC) with similar efficiency (Fig. 2J, S3A), which is inconsistence with plaque assays, indicating the difference between *in vivo* and *in vitro* restrictions. We also expressed mzaD mutants, D187A and P297A (Fig. S3B). At the high reaction concentrations, both of the mutants show decreased efficiency in digestion phage DNAs compared to the wild-type mzaD whereas the nuclease activities of mutants were almost abolished at low protein concentration (0.5 μM) (Fig. S3C). Compared to T4 (ghmC), both T4 (C) and T4 (hmC) genomic DNAs are more sensitive to the mzaD mutants (Fig. S3C). These results are consistent with plaque assays that both mutants have no activity against T4 (ghmC) but have lower activity against T4 (C) and T4 (hmC) compared to the wild-type system (Fig. 2E), indicating mzaABCDE directly targets T4 genomic DNA through its nuclease activity.

**Fig. 3.**
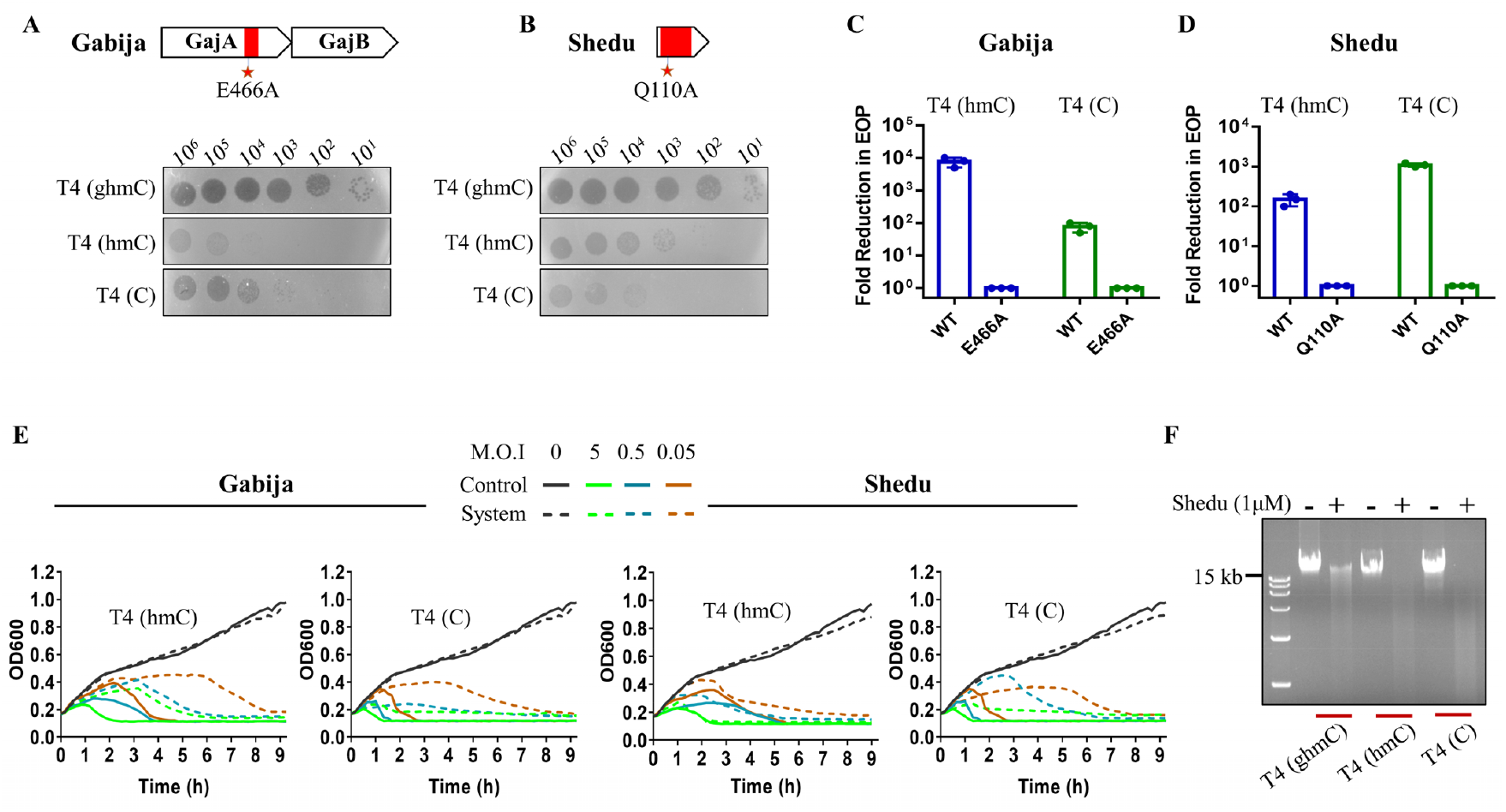
Glucosylation of hmC in phage T4 genome abolishes the defense activities of Gabija and Shedu. Genetic compositions and defense activities of *E. coli* Gabija **(A)** and Shedu **(B)** against T4 (ghmC), T4 (hmC), and T4 (C). The nuclease domains were highlighted with red, and the putative active sites in nuclease domain were indicated with red starts. The representative results of plaque assays were shown. The antiphage activities of wild-type and mutated Gabija **(C)** and Shedu **(D)** against phages T4 (hmC) and T4 (C) were presented as EOP. Data were represented as mean ± S.D. **(E)** Growth curves of *E. coli* DH10B cells expressing Gabija or Shedu after infection with T4 (hmC) and T4 (C) at different MOI. **(F)** The representative results of *in vitro* nuclease assay showing the nuclease activity of purified Shedu protein on genomic DNAs of phages T4 (ghmC), T4 (hmC), and T4 (C).

### Glucosylation of hmC in phage T4 genome abolishes the defense activities of Gabija and Shedu

We found Gabija and Shedu can protect *E. coli* cells against both T4 (C) and T4 (hmC) phages but their defense activities were completely blocked by wild-type phage T4 where hmCs in genomic DNA were further glucosylated (Fig. 3A, B). The antiphage activity of Shedu significantly decreased when cytosines in T4 genomic DNA were replaced with methylcytosines, with 10^3^- and 10^2^-fold protection against cell lysis for phages T4 (C) and T4 (hmC) respectively (Fig. 3B). Interestingly, Gabija showed higher defense efficiency against T4 (hmC) compared to T4 (C) (10^3^-fold versus 10-fold reduction, Fig. 3A).

Mutations on the putative active sites or the conserved amino acids in the nuclease domain showed that the E466A mutation completely abolished the defense activity of Gabija against both T4 (C) and T4 (hmC) phages (Fig. 3 A and C). Similarly, mutations at site Q110A also completely abolished the defense activity of Shedu (Fig. 3 B and D). These results indicated that the antiphage activities of Gabija and Shedu depend on their nuclease activities. The growth kinetics of phage-infected *E. coli* cells expressing Gabija or Shedu did not show any evidences for the defense system-mediated growth arrest no matter how many phages were used for infections (Fig. 3E). These results indicated that both Gabija and Shedu systems restrict T4 (C) and T4 (hmC) through directly targeting phages rather than inducing abortive infection, and ghmC-modification confers T4 resistances against these systems.

To further determine the resistance of different modified T4 genomic DNAs against nucleases of Gabija and Shedu, we expressed the recombinant GajA and Shedu using *E. coli* BL21 cells. However, due to the solubility issue of GajA protein, we only got Shedu protein (Fig. S4A). Nuclease assay showed that the recombinant Shedu efficiently digested genomic DNAs of both T4 (C) and T4 (hmC) phages, whereas ghmC-modified genomic DNA is partially resistant against Shedu nuclease (Fig. 3F and S4B). These results are consistent with the plaque assays that Shedu system has defense activity against phages T4 (C) and T4 (hmC) but not against wild-type T4, indicating the defense mechanism of Shedu against T4 mutants via degradation of phage genomic DNA.

### The Restriction-like defense system specifically targets phage T4 (hmC)

The Restriction-like defense system was firstly identified in 2020, which is composed of 4 proteins containing nuclease domain, ATPase domain, methylase domain, helicase domain, and several uncharacterized domains (Fig. 4A) [25]. It was shown that this system is active against phages P1, λ, and M13 with variable efficiency but inactive against phages T2, T3, T4, T5, and T7 [25]. Our plaque assays also confirmed that the Restriction-like system lacks the defense activity against wild-type T4 (Fig. 4B). However, we found it can specifically protect *E. coli* cells against phage T4 (hmC), with 10^2^-fold protection against cell lysis, but not against phage T4 (C) (Fig. 4B). Mutations on the putative active sites G132A, K135A, H233A, and T269A in the nuclease domain did not affect the antiphage activity of Restriction-like system against phageT4 (hmC). Interestingly, mutations on sites Q109A and D230A even increased the defense activity (Fig. 4C). These results indicated that anti-T4(hmC) activity of Restriction-like defense system does not depend on nuclease activity. The growth kinetics of phage-infected *E. coli* cells expressing Restriction-like system did not show any evidences for the growth arrest no matter how many phages were used (Fig. 4D). However, *E. coli* cells kept growing within the first 3h after infection with 5 MOI of T4 (hmC) (Fig. 4D), indicating a delay in assembly of progeny phages and, therefore, the subsequent cell lysis.

**Fig. 4.**
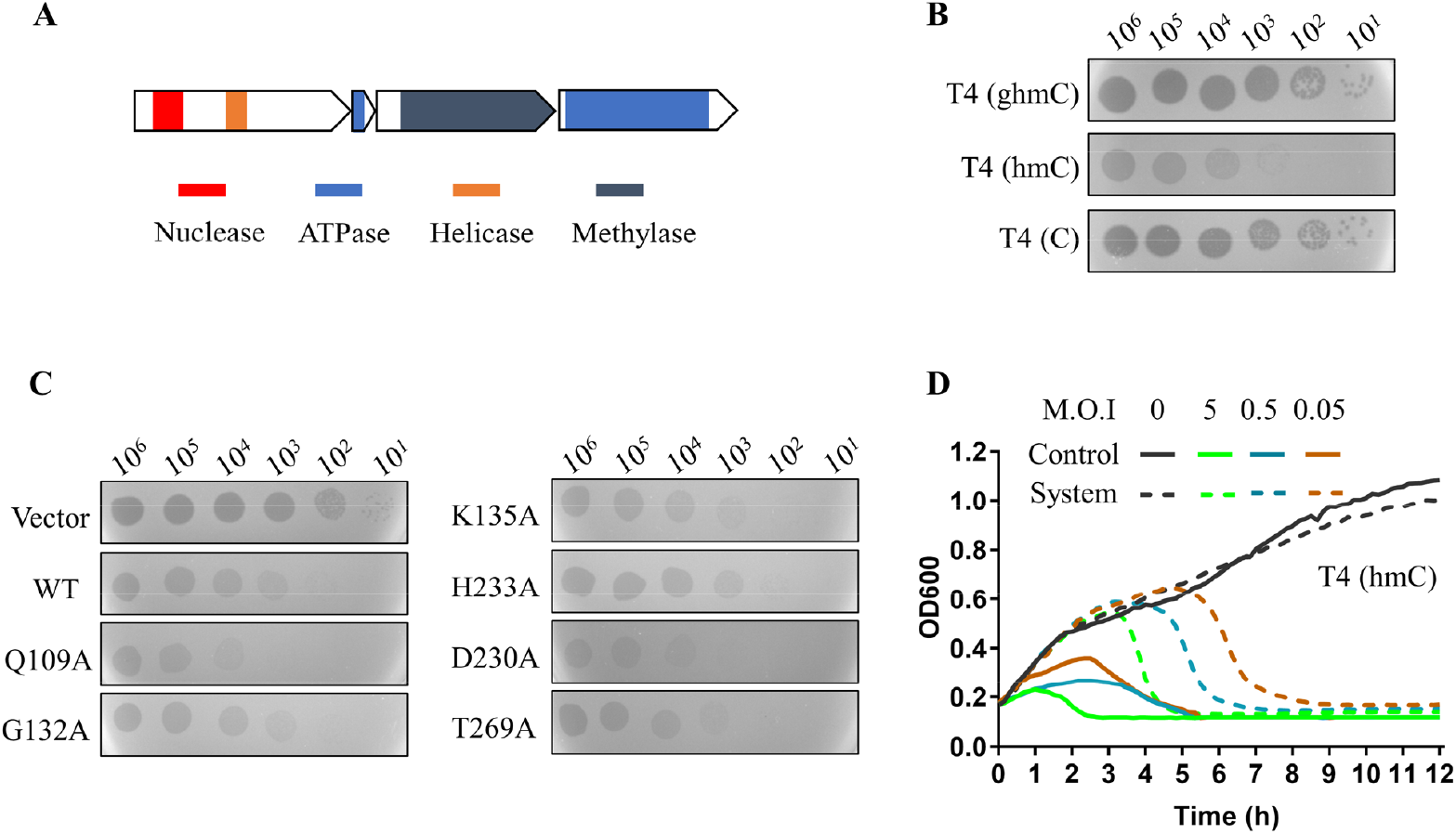
The Restriction-like defense system specifically targets phage T4 (hmC). **(A)** Cartoon shows the genetic composition and protein domain of Restriction-like system. The main functional domains were indicated with different colors. **(B)** The representative results of plaque assays showing the defense activity of Restriction-like system against T4 (ghmC), T4 (hmC), and T4 (C). The amount of phages (10-10^6^ pfu) used for plaque assays were indicated on top of each panel. **(C)** Phage plaque assays on *E. coli* DH10B cells expressing wild-type or mutated Restriction-like systems. *E. coli* cells containing empty vector were used as controls. The mutation sites, Q109A, G132A, K135A, H233A, D230A, T269A, in nuclease domain were indicated. **(D)** Growth curves of *E. coli* DH10B cells in the presence (dashed lines) and absence (solid lines) of the Restriction-like system after infection with T4 (hmC) at MOI of 0, 0.05, 0.5, and 5.

### Druantia and qatABCD system confer protection only against phage T4 (C)

The Druantia defense system was discovered in 2018, which is characterized by a very large protein DruE (∼2,000 amino acids long and contains an unannotated domain DUF1998) accompanied by 1-4 accessory proteins [23]. Three types of Druantia systems (I-III) were reported based on the number of accessory proteins, however, only types I and III were found in *E. coli* in our current study with type III as the dominant system (639 of total 667, Fig. S5). The Type I Druantia system was shown to have defense activity against wild-type phage T4, but its DruE does not contain a nuclease domain [23]. We, therefore, cloned the Type III Druantia, which contains only 1 accessory protein and is the most abundant nuclease-containing system in *E. coli* (Fig. 1C). We found that the type III Druantia system has no defense activity against wild-type T4 and T4 (hmC), but is highly active against phage T4 (C), with 10^6^-fold protection against cell lysis (Fig. 5A). These results revealed a defense and counter-defense mechanism between *E. coli* and phage T4, namely, phage resists type III Druantia via ghmC-modification of its genome whereas *E. coli* cells employ type I Druantia to overcome this resistance. Another nuclease-containing system that only targets phage T4 (C) rather than wild-type T4 and T4 (hmC) is qatABCD (Fig. 5B), which was identified in 2020 as a defense system against phages P1, T3, and λ but not against phages T2, T4, T5, and T7 [25]. However, the activity of qatABCD against T4 (C) is quite low compared to the type III Druantia system (10-fold versus 10^6^-fold reduction, Fig. 5A, B).

**Fig. 5.**
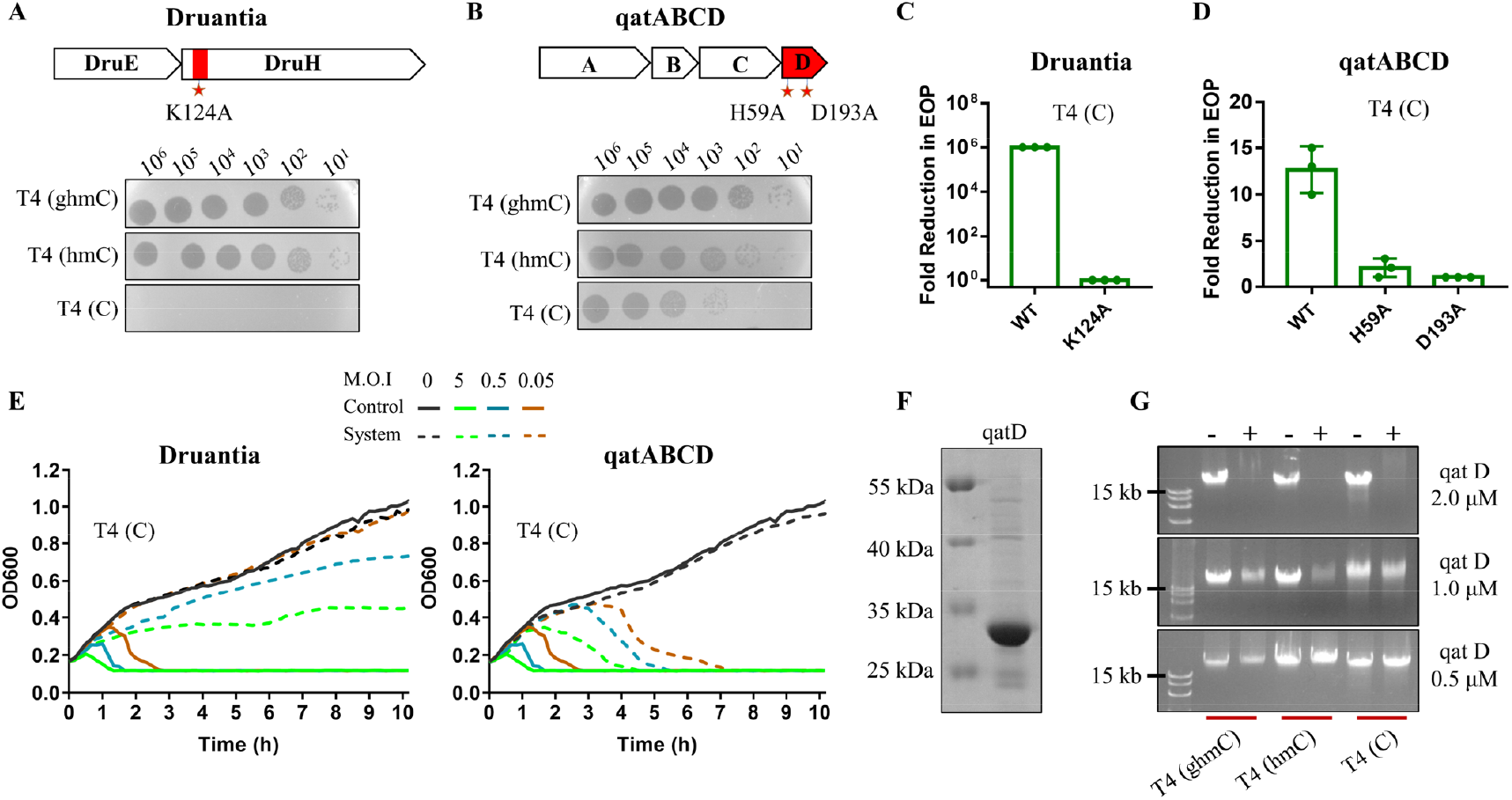
Druantia and qatABCD system confer protection only against phage T4 (C). The genetic composition and defense activities of *E. coli* Druantia **(A)** and qatABCD **(B)** against T4 (ghmC), T4 (hmC), and T4 (C). Genetic composition of each system, nuclease domains (highlighted with red), and the putative nuclease active sites (red starts) were shown on top of each panel. The representative results of plaque assays were shown at the bottom. The defense activities of wild-type and mutants of Druantia **(C)** and qatABCD **(D)** against phage T4 (C) were presented as EOP. Data were represented as mean ± S.D of three independent assays. **(E)** Growth curves of *E. coli* DH10B cells expressing Druantia or qatABCD after infection with phage T4 (C) at MOI of 0, 0.05, 0.5, and 5. The qatD protein of qatABCD system was purified **(F)**, and its nuclease activity on genomic DNAs of phages T4 (ghmC), T4 (hmC), and T4 (C) was determined by incubating 350 ng DNA with different amount of qatD protein **(G)**.

Mutations on the putative active sites in the nuclease domain showed that the K124A mutation completely abolished the defense activity of type III Druantia against phage T4 (C) (Fig. 5A, C). Similarly, mutations at sites H59A and D193A completely abolished the defense activity of qatABCD against T4 (C) (Fig. 5B, D). These results indicated that the nuclease activity is necessary for type III Druantia and qatABCD to defend *E. coli* cells against phage T4 (C). The growth kinetics assays of *E. coli* cells expressing type III Druantia and qatABCD also supported the concept that these defense systems directly target phage T4 (C) rather than inducing abortive infection (Fig. 5E).

To determine whether these systems directly degrade phage genomic DNAs, we expressed nuclease domain-containing proteins, DruE and qatD, from Druantia and qatABCD respectively. However, we were only able to purify the recombinant qatD (Fig. 5F). Interestingly, we found qatD can equally degrade genomic DNAs of wild-type and T4 mutants at a high reaction concentration (2.0 μM), and the nuclease activity is dose-dependent (Fig. 5G). These results indicated that the qatABCD system might restrict T4 phage via direct degradation of phage genome. However, more experiments are need to reveal how ghmC- and hmC-modified T4 DNAs resist qatD degradation. One explanation for the inconsistency of nuclease and plaque assays is the reaction condition difference between *in vivo* and *in vitro*.

### Six nuclease-containing systems lack the defense activities against phage T4

We found six nuclease-containing systems recently identified, namely Zorya, hhe, ppl, AVAST type 4, Retron Ec78, and Retron Ec67-like, lack the defense activities against wild-type phage T4 and the mutants (Fig. 6A). There are two types of Zorya systems, both of which contain a nuclease domain. We were able to clone type II Zorya that contains an HNH-nuclease domain from our *E. coli* collection. Plaque assay showed type II Zorya has defense activity against phage T7, and mutation on the putative active site in the nuclease domain significantly reduced the defense activity (Fig. 6A, B). However, type II Zorya is not active against phage T4 regardless of the genome modifications (Fig. 6A), indicating T4 possesses other counter-defense mechanisms rather than genome modifications or type II Zorya lacks the ability to recognize T4 infection.

**Fig. 6.**
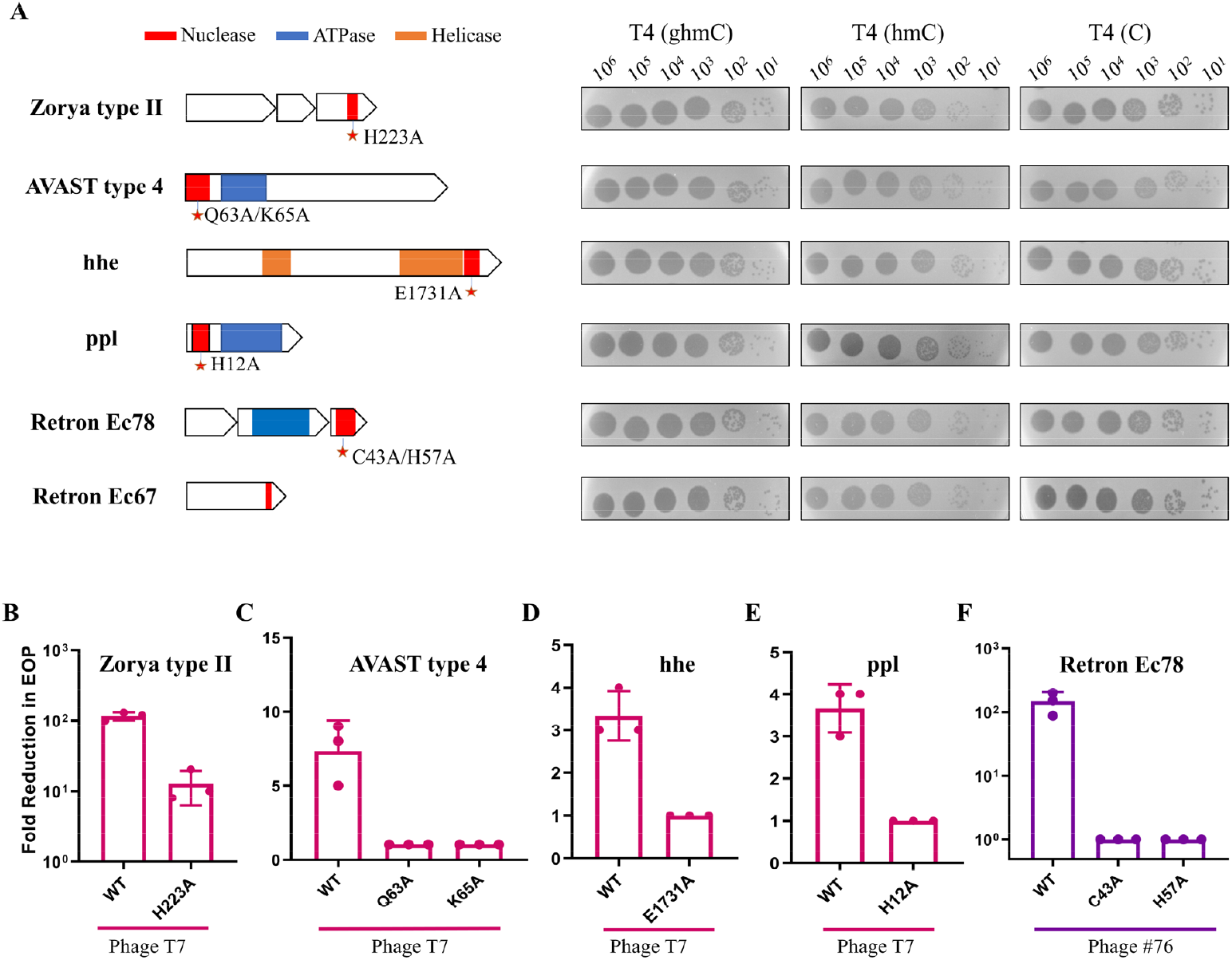
*E. coli* Zorya, hhe, ppl, AVAST type 4, and Retron Ec78 systems lack the defense activities against phage T4. **(A)** The genetic composition and defense activities of each system against phages T4 (ghmC), T4 (hmC), and T4 (C). The genetic compositions, main functional domains, and the putative nuclease active sites (red starts) were shown on left side. The antiphage activities were determined by plaque assays and representative results were shown. **(B)** Antiphage activities of wild-type and mutants of each system against phage T7 or phage #76. Data were represented as mean ± S.D of three independent phage plating assays.

AVAST type 4, which contains an Mrr-like nuclease domain, has defense activity against phages T3, T7, and ΦV-1[25]. In our study, we also found it can protect *E. coli* cells against phage T7 but with very low defense activity, which was abolished when the putative active site in the nuclease domain was mutated (Fig. 6A, C). Similarly, we did not find the defense activity of AVAST type 4 against phages T4 and the mutants (Fig. 6A).

The hhe system contains a Vsr (Very short patch repair) endonuclease domain and is active against phages λ, T3, T7, and ΦV-1 [25]. The ppl system contains a nuclease domain at its NH-terminus and was also shown active against phages λ, T3, T7, and ΦV-1 [25]. In our study, we found both systems have very weak defense activities against phage T7, and mutations on the sites E1731A of hhe and H12A of ppl abolished antiphage activities of each system respectively (Fig. 6A, D and E). These results are consistent with previous studies [25]. Similar as type II Zorya and type 4 AVAST, both hhe and ppl systems are incapable to restrict T4 infection no matter wild-type, T4 (hmC), or T4 (C) (Fig. 6A).

Retron Ec78 has an HNH-nuclease domain and was shown to have a defense activity against phage T5, which was not affected when the putative active site (H57A) in the nuclease domain was mutated [25]. However, our phage T5 stock, which showed sensitivity against Rectron Ec86 in our recent study, can completely tolerate Ec78. The defense activity of Ec78 was confirmed using phage #76, which was completely abolished when the nuclease active sites C43A and H57A were mutated (Fig. 6A, F). However, Ec78 didn’t show defense activity against both wild-type T4 and the mutants. In addition, we identified a retron system in *E. coli* that contains a predicted HNH-nuclease domain and has similar genetic organization as retron Ec67. However, this Ec67-like system did show defense activity against T4 (ghmC), T4 (hmC), T4 (C), T5, and all other 111 phages (Fig. 6A).

Previous studies indicated that capsid-targeted internal protein 1 (IPI) of T4, which was packaged in the mature phage particles and injected into host cells during infection, can protect T4 genomic DNAs from type IV R-M nuclease [30]. To determine whether IPI confers T4 phages resistances against these six nuclease-containing systems, we deleted *IPI* gene from the wild-type T4 genome (Fig. S6A). Plaque assay showed that the T4 *ΔIPI* mutant is still resistant against these nuclease-containing systems (Fig. S6B), indicating that T4 phage might employ other mechanisms to invade these defense systems.

## DISCUSSION

Phage T4 is a widely used model to explore the mechanisms of phages to resist nucleases of *E. coli* R-M systems, which had revealed the ghmC modification as one of the most powerful covalent modifications that makes T4 DNA resistant against most of types I-III R-M systems. It leads to the concept that hypermodification of genomic DNA could be a key strategy of phages to counteract bacterial defense systems. Recently, many new defense systems that contain nuclease domains were identified from a variety of bacteria and archaea, however, many of them have not been identified or validated in *E. coli*. Here, we systematically analyzed nuclease-containing defense systems employed by *E. coli* and the corresponding counter-defense activities of T4 ghmC modification.

By analyzing 2,289 genome sequences available in NCBI, we found *E. coli* employed at least 15 of 20 recently identified nuclease-containing systems to protect themselves against phages. We were able to clone 14 systems from our *E. coli* collections and test their protection activities. Particularly, Gabija, Septu, Shedu, mzaABCDE, and type III Druantia were firstly identified and validated in *E. coli*. The Gabija system has broad antiphage activity (97 of 111 phages), whereas Septu and Shedu inhibit only 10-11 phages. Druantia systems are divided into three types (I-III), and only type I has been validated in *E. coli*. We only found types I and III in the 2,289 *E. coli* genomes, with type III as the dominant type (639 of total 667), which is also the most abundant nuclease-containing system in *E. coli*. We found that the type III Druantia also has broad antiphage activity against 70 out of the 111 tested phages, indicating that it is a key defense system of *E. coli*. Another system worth mentioning in our studies is SspBCDE, the homology of *V. cyclitrophicus* SspABCD–SspE that restricts phages through introducing nicks to the genomic DNAs by SspE. The nickase activity of SspE is stimulated by PT modification, which is carried out by SspA, B, C, and D. Both PT modification and SspE of *V. cyclitrophicus* are necessary for its antiphage activity [24]. However, we found SspBCDE, which is defective in PT modification due to the lack of SspA, restricts the infection of wild-type T4 phage as well as two mutants, indicating the different defense mechanism of *E. coli* SspBCDE system.

Like most phages, T4 genomic DNAs are injected into *E. coli* cells during infection to initiate genome replication, protein expression, and subsequent virion assembly. Therefore, its genomic DNA is considered as a main target for *E. coli* to restrict infections, and it is expected that *E. coli* cells employ nuclease-containing systems to degrade phage DNAs whereas phages evolve the counter-defense strategies via modifying their genomic DNAs. Indeed, we found the ghmC-modification of phage T4 genome can completely block the defense activities of Gabija, Shedu, Restriction-like, type III Druantia, and qatABCD. The antiphage activities of the last two defense systems can also be counteracted by hmC-modification (Fig. 3-5). Interestingly, Gabija system is more effective to restrict T4 (hmC) than T4 (C), and the Restriction-like system is only able to restrict T4 (hmC) (Fig. 3). However, the antiphage activities of Septu, SspBCDE, and mzaABCDE were not abolished by ghmC- or hmC-modifications of T4 phage. In particular, Septu system showed similar activities against phages T4 (ghmC), T4 (hmC), and T4 (C). Although SspBCDE and mzaABCDE can restrict both wild-type T4 and two mutants, their antiphage activities heavily depends on the extent of cytosine modifications in T4 genome (Fig. 2). In addition, there are six nuclease-containing systems that cannot protect *E. coli* cells against phage T4 no matter the cytosines are modified or not. Taken together, we showed covalent modifications of cytosines confers T4 phage variable resistance against some nuclease-containing systems but *E. coli* can use other nuclease-containing systems to target T4 phage with different modifications, and T4 phage might have other mechanisms in addition to DNA modifications to counteract nuclease-containing systems.

Mechanism studies indicated that different nuclease-containing systems use different mechanisms to restrict T4 phage infection, and even the same defense system might adopt different approaches to restrict T4 phages with different modifications. For instance, *E. coli* cells employ Shedu system to degrade T4 genome DNAs through its nuclease activity (Fig. 3) whereas the nuclease activity of Restriction-like system is not necessary for its antiphage ability and mutations on nuclease active sites even enhance its defense activity (Fig. 4). A single point mutation in the nickase catalytic center of SspE completely abolished the defense activities of SspBCDE against T4 (C) and T4 (ghmC), but did not affect the activity against T4 (hmC) (Fig. 2). Similarly, growth arrest was only observed when SspBCDE-containing *E. coli* cells were infected by phage T4 (hmC) but not T4 (ghmC) and T4 (C), indicating different a defense mechanism of SspBCDE against T4 (hmC).

In conclusion, we showed *E. coli* employs diversified nuclease-containing systems to protect themselves against phage infection. The ghmC modification is a critical counter-defense mechanism by phage T4 that makes it resistant against multiple but not all nuclease-based defense systems. Divergent antiphage mechanisms of nuclease-containing systems, other than directly degrading phage DNA, help *E. coli* overcome ghmC modification of phage T4 and provide *E. coli* with multidimensional defense strategies. However, T4 phage have evolved other unrevealed counter defense mechanisms to establish the ecological balance and the greatest diversity of both bacteria and phage in the biosphere.

## MATERIAL AND METHODS

### Bacteria and phages

*E. coli* strains DH5α (*hsdR17(rK–mK+) sup*^*2*^), DH10B (F^−^ *mcrA Δ(mrr-hsdRMS-mcrBC)*), MG1655 (F^−^ λ^−^ *rph-1*), B834 (*hsdR*_*B*_ *hsdM*_*B*_ *met thi sup*^0^), B40 (sup^1^), and BL21 (DE3) were used in this study as described below. *E. coli* DH5α was used for plasmid construction, and B834 was used for construction of T4 mutants, T4 (hmC) and T4 (C), as described below. *E. coli* MG1655 was initially used to determine the antiphage activities of all the nuclease-containing defense systems. The defense activities of nuclease-containing systems against T4 (ghmC), T4 (hmC), and T4 (C) were carried out with *E. coli* DH10B. A collection of 816 *E. coli* strains from our laboratory stocks was used to amplify nuclease-containing defense systems. Phages T4, T5, and T7 were from our laboratory stocks. A collection of 111 phages, which we recently isolated with *E. coli* MG1655 as host cells, was used to determine the anti-defense effects of 5 potential defense systems in *E. coli*.

### Identification of the nuclease-containing defense systems in *E. coli*

All complete *E. coli* genomes (2,289 strains as of February 2021) available at NCBI were downloaded, and the protein coding sequences were analyzed using Prodigal (release 2.6.3) [31]. Protein domains were annotated using HMMER version 33.0 [27]. The conserved domains of all 20 nuclease-containing defense systems identified as of February 2021 (Table S1) were collected. The *E. coli* protein sequences were screened using blast+ (release 2.11.0) for the potential defense systems that share similar domain structures with reported nuclease-containing systems. A collection of 816 *E. coli* strains from our laboratory stocks was used to amplify identified defense systems using primers listed in Table S7. We were able to amplify 14 defense systems (Table S5) from our *E. coli* colletion and clone to pSEC1 vector (Table S8), which includes a kanamycin-resistent gene and the p15A origin of replication. The generated plasmids were confirmed by sequencing.

### Generation of T4 mutants

T4 mutants, T4 (hmC) and T4 (C), were generated using CRISPR-Cas12 phage genome editing technology [32]. Construction of the pLbCas12a plasmids, which express different crRNAs targeting different genes with T4 genome, were carried out as described in our previous study [14]. Briefly, the spacer fragments were cloned into the pLbCas12a under the control of J23100 promoter to express the corresponding crRNA by Gibson assembly of spacer fragments and EcoRI/XhoI-linearized pLbCas12a vector. Restriction efficiency of each pLbCas12a-spacers plasmid on T4 phages infection was determined by plaque assays and showed in Table S6. The donor plasmids were constructed by cloning the fragments containing the gene of interest flaking with two homologous arms into XbaI/EcoRI- or KpnI/EcoRI-linearized vector. The primers are available upon request. The CRISPR-LbCas12a plasmid and the corresponding donor plasmid were co-transformed into *E. coli* DH10B or B834, which were used for phage genome editing. *E. coli* cells transformed with the CRISPR-Cas12a plasmid only was used as a control. About 10^8^ PFU of phages were incubated with 200μL of log-phase *E. coli* cells at 37°C for 7 min. After adding 3 milliliters of 0.75% LB top agar containing appropriate antibiotics, the mixture was poured onto a TSA-agar plate and incubated at 37°C overnight. The recombinant T4 phages were confirmed using PCR and sequencing of single plaques.

### Phage plaque assay

Phage plaque assays were carried out as described previously [21]. Briefly, 500µL *E. coli* fresh culture was mixture with 8 mL top agar (30g/L TSB, 7.5g/L agar, and 50 µg/mL Kanamycin) and poured onto a TSA-agar plate. Then, 100 μL of 10-fold serially diluted phages were dropped on the plate, and phage plaques were counted after overnight incubation at 37 °C. Fold reduction in efficiency of plating (EOP) was calculated by dividing the input pfu by the number of plaques produced from infection of *E. coli* expressing the defense system, and expressed as mean ± S.D. of three independent assays. For the defense spectrum experiments, the plasmid expressing specific defense system was transformed into *E. coli* MG1655 cells, which were then infected with 10-fold serially diluted phages. Totally, 111 different phages isolated from different places in China were used. *E. coli* MG1655 cells transformed with an empty vector, pSEC1, were used as controls. For T4 phage and its mutants infection experiments, *E. coli* DH10B cells were used instead of MG1655, in which phages T4 (hmc) and T4 (C) cannot propagate.

### Phage-infection dynamics in liquid medium

An overnight culture of *E. coli* DH10B containing the plasmid that expresses specific defense system was inoculated into fresh LB medium supplemented with 50 µg/mL Kanamycin and incubated at 37 °C with a shaking speed of 200 rpm. *E. coli* cells containing the empty plasmid were used as controls. When the cell density reaches OD600 of 0.3, 180 µl of cultures were dispensed into each microwell in a 100-well plate containing 20 µl of 10-fold serially diluted phages at a MOI of 5, 0.5, and 0.05, respectively. The plates were instantly placed on a automated growth curve analysis system (Oy Growth Curves Ab Ltd Bioscreen C). The *E. coli* cells were kept at 37°C with continuous shaking, and the OD600 was measured every 10 min for a total duration of 12h.

### Expression and purification of protein

Genes containing nuclease domains were amplified by PCR and cloned into BamHI/XhoI-linearized pET-28a vector using T4 DNA ligase. The generated plasmids were confirmed by sequencing and then transformed into *E. coli* BL21 (DE3). Protein expression was induced with 1mM isopropyl-β-D-thiogalactoside (IPTG) at 15°C for 16 hours when the OD600 of culture reached 0.6. Protein was purified as described previously [33]. Briefly, *E. coli* cells were pelleted by centrifugation at 8000g at 4 °C for 10 min and lysed at 12000 psi using a French press. After centrifugation at 30000g at 4 °C for 30 min, the supernatant was collected and filtered through 0.22μm filters. The protein was purified affinity chromatography using Ni-NTA column (Yeasen, Wuhan, China) and concentrated using centrifugal ultrafiltration (Millipore). The concentration of recombinant protein was determined with bovine serum albumin (BSA) as a standard.

### Phage propagation and genomic DNA extraction

Wild-type T4 and two mutants, T4 (hmC) and T4 (C), were purified as previously described [34, 35]. Briefly, log phase *E. coli* DH10B cells (∼2×10^8^ cells per ml) grown on LB medium were infected with phages at MOI of 0.1. After 5 h incubation at 200 rpm at 37 °C, the *E. coli* cells were collected by centrifugation at 30,000 g for 50 min. The precipitates were resuspended with 10 ml of Pi-Mg buffer (26 mM Na_2_HPO_4_, 68 mM NaCl, 22 mM KH_2_PO_4_, 1 mM MgSO_4_, pH 7.5) containing 10 μg/mL DNase I and incubated at 37 °C for 1h. Phages genomic DNAs were isolated with the phenol-chloroform-isoamyl alcohol extraction. Phage pellet was firstly digested with 100 µl 10% SDS and 100 µl 0.5 mol /L EDTA in a 65 °C water bath for 30 min. A mixture of phenol:chloroform:isoamyl alcohol (volume ratio of 25: 24:1, 10ml) was added and inverted several times until the emulsion formed. The supernatant was collected after centrifugation at 10,000 g at 4 °C for 10 min and subjucted to a second-round extraction. The genomic DNAs were precipitated by adding isopropanol. After centrifugation at 10,000 g at 4 °C for 10 min, the genomic DNA pellet was washed twice with 1 mL precooled 70% ethanol and dissolved in 200 μ L deionized water.

### Nuclease assays

The modifications of cytosines in genomic DNAs of T4 (ghmC), T4 (hmC), and T4 (C) were determined by digerstion with QuickCut™ EcoR V (Takara), MspJI (NEB), and AluI (NEB), respectively. The nuclease activities of recombinant proteins were determined by incubation of phage T4 genomic DNAs (350 ng) with different amount of recombinant proteins (2μM, 1μM, and 0.5μM) in a final volume of 10 µl in DNA cleavage buffer (1mM NTP Mix (Beyotime), 20 mM Tris–HCl (pH 9.0), 0.1 mg/ml BSA, and 5mM MgCl_2_). After 5 min incubation at 37 °C, the reactions were stopped by adding 2µl of 6× loading dye (NEB) containing 20 mM EDTA and analyzed via agarose gel electrophoresis.

## ACKNOWLEDGEMENTS

Our researches are supported by National Natural Science Foundation of China (Grant No. 32170094) and Fundamental Research Funds for the Central Universities (Program No. 2662022DKYJ003).

