## Supplementary for "The complex roles of genomic DNA modifications of bacteriophage T4 in resistance to nuclease-based defense systems of *E. coli*"

^b^ Hongshan Lab, Wuhan, Hubei 430070, China.

^c^ Key Laboratory of Development of Veterinary Diagnostic Products of Ministry of Agriculture, Huazhong Agricultural University, Wuhan, Hubei 430070, China

^d^ Bacteriophage Medical Research Center, Department of Biology, The Catholic University of America, Washington, DC 20064, USA

**^#^** These authors contributed equally to this work.

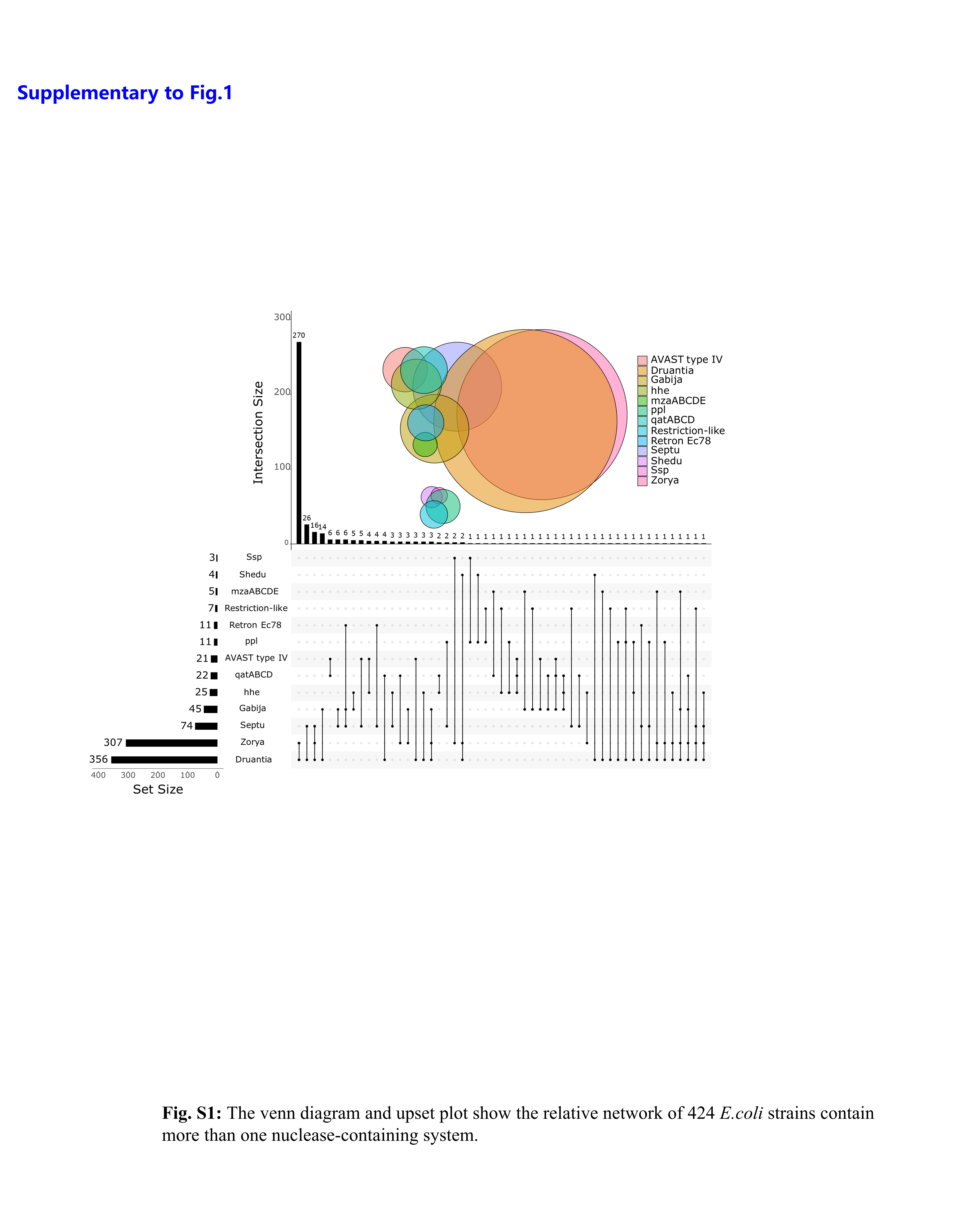
**Fig. S1**

**Fig. S1.** The venn diagram and upset plot show the relative network of 424 *E. coli* strains contain more than one nuclease-containing system.

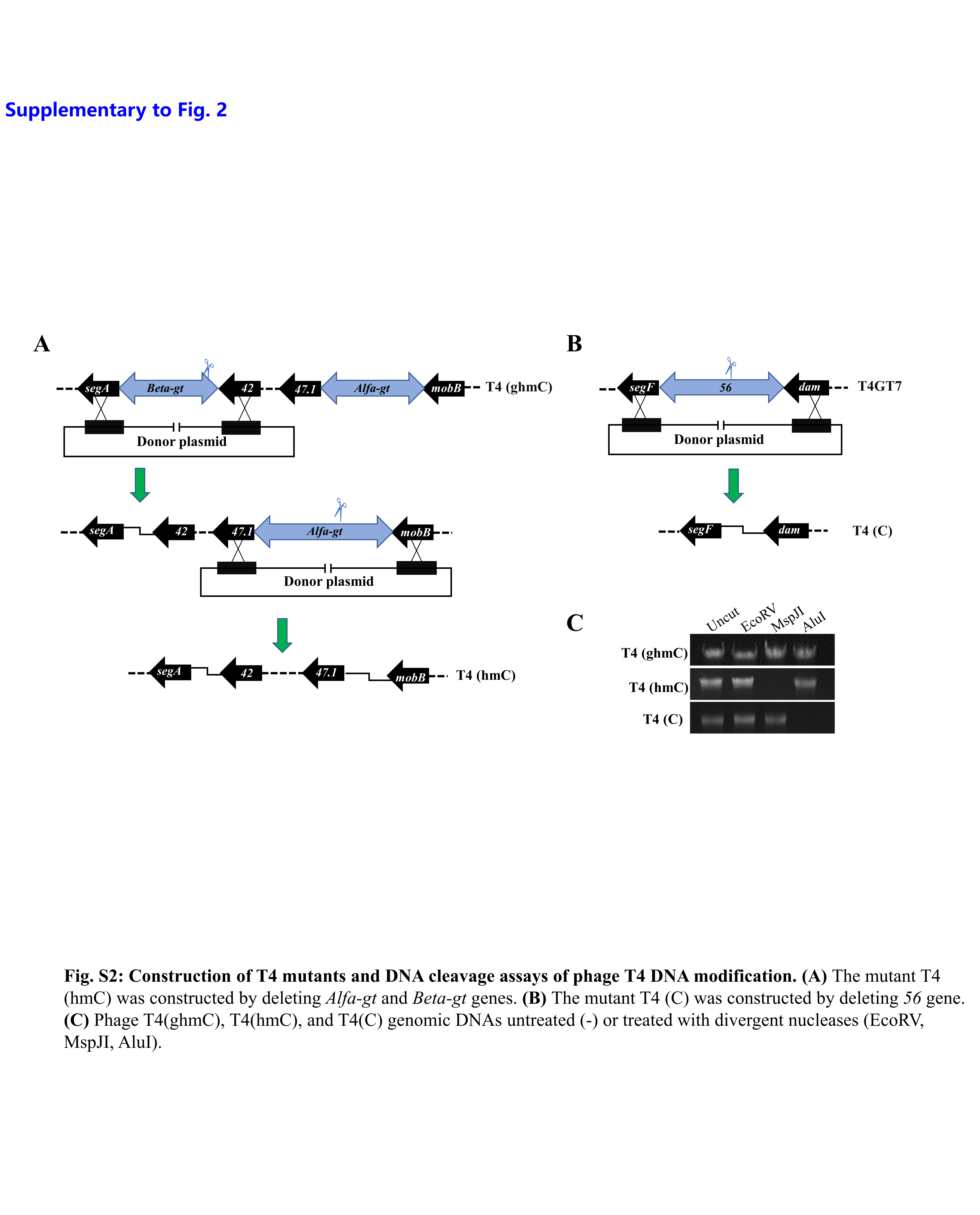
**Fig. S2**

**Fig. S2. Construction of T4 mutants and DNA cleavage assays of phage T4 with different DNA modifications. (A)** The mutant T4 (hmC) was constructed by deleting *Alfa-gt* and *Beta-gt* genes. **(B)** The mutant T4 (C) was constructed by deleting *56* gene. **(C)** Phage T4(ghmC), T4(hmC), and T4(C) genomic DNAs were treated or untreated (Uncut) with divergent nucleases (EcoRV, MspJI, and AluI).

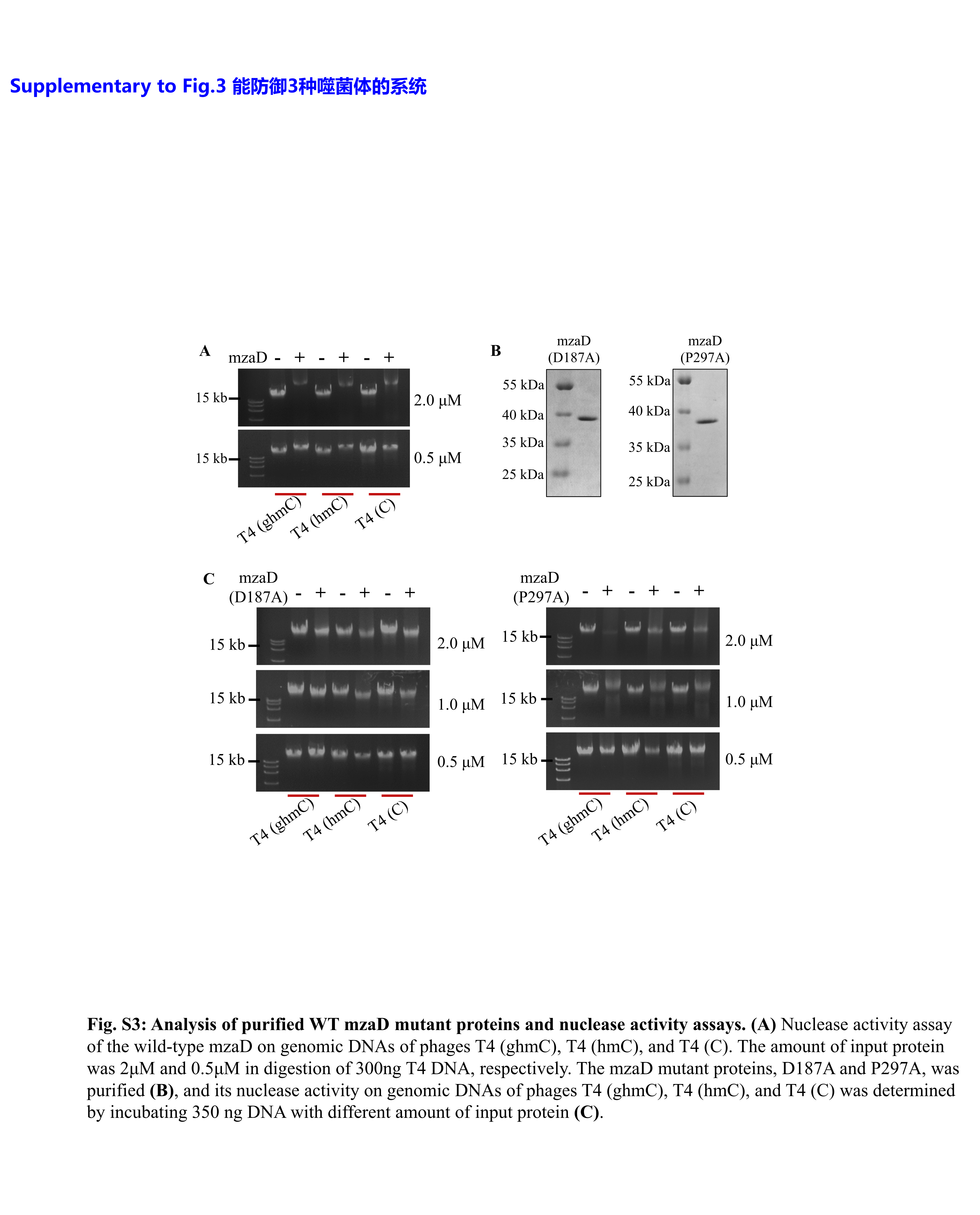
**Fig. S3**

**Fig. S3. Nuclease activities of wild-type and mutants mzaD proteins. (A)** Nuclease activity assay of the wild-type mzaD on genomic DNAs of phages T4 (ghmC), T4 (hmC), and T4 (C). Three hundred and fifty ng T4 DNA was incubated with 2μM or 0.5μM mzaD protein. The mzaD mutant proteins, D187A and P297A, were purified **(B)**, and its nuclease activity on genomic DNAs of phages T4 (ghmC), T4 (hmC), and T4 (C) was determined by incubating 350 ng DNA with different amount of recombinant proteins **(C)**.

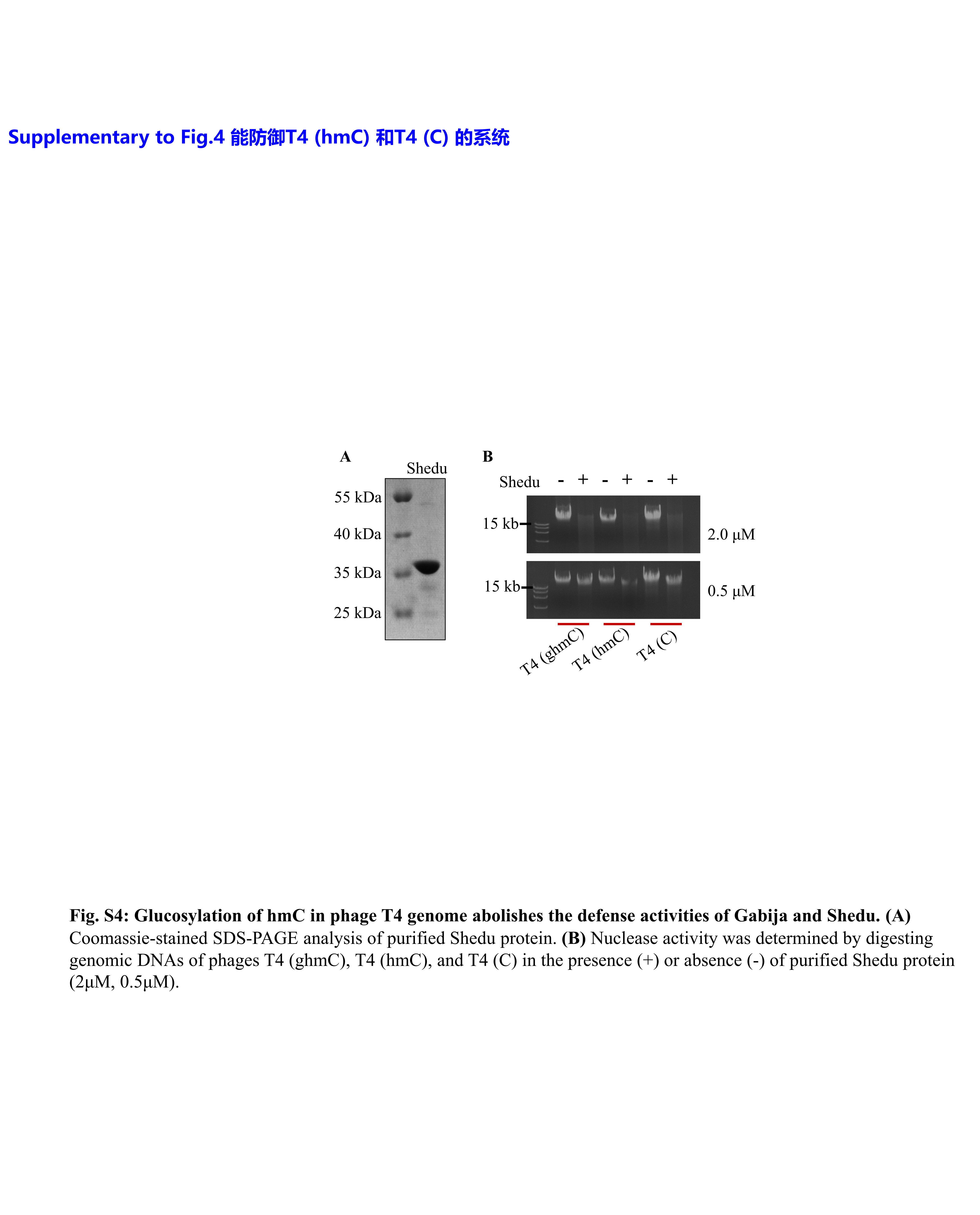
**Fig. S4**

**Fig. S4. Glucosylation of hmC in phage T4 genome abolishes the defense activities of Shedu. (A)** SDS-PAGE analysis of purified Shedu protein. **(B)** Nuclease activity was determined by digesting genomic DNAs of phages T4 (ghmC), T4 (hmC), and T4 (C) in the presence (+) or absence (-) of recombinant Shedu protein (2μM or 0.5μM as indicated in the figure).

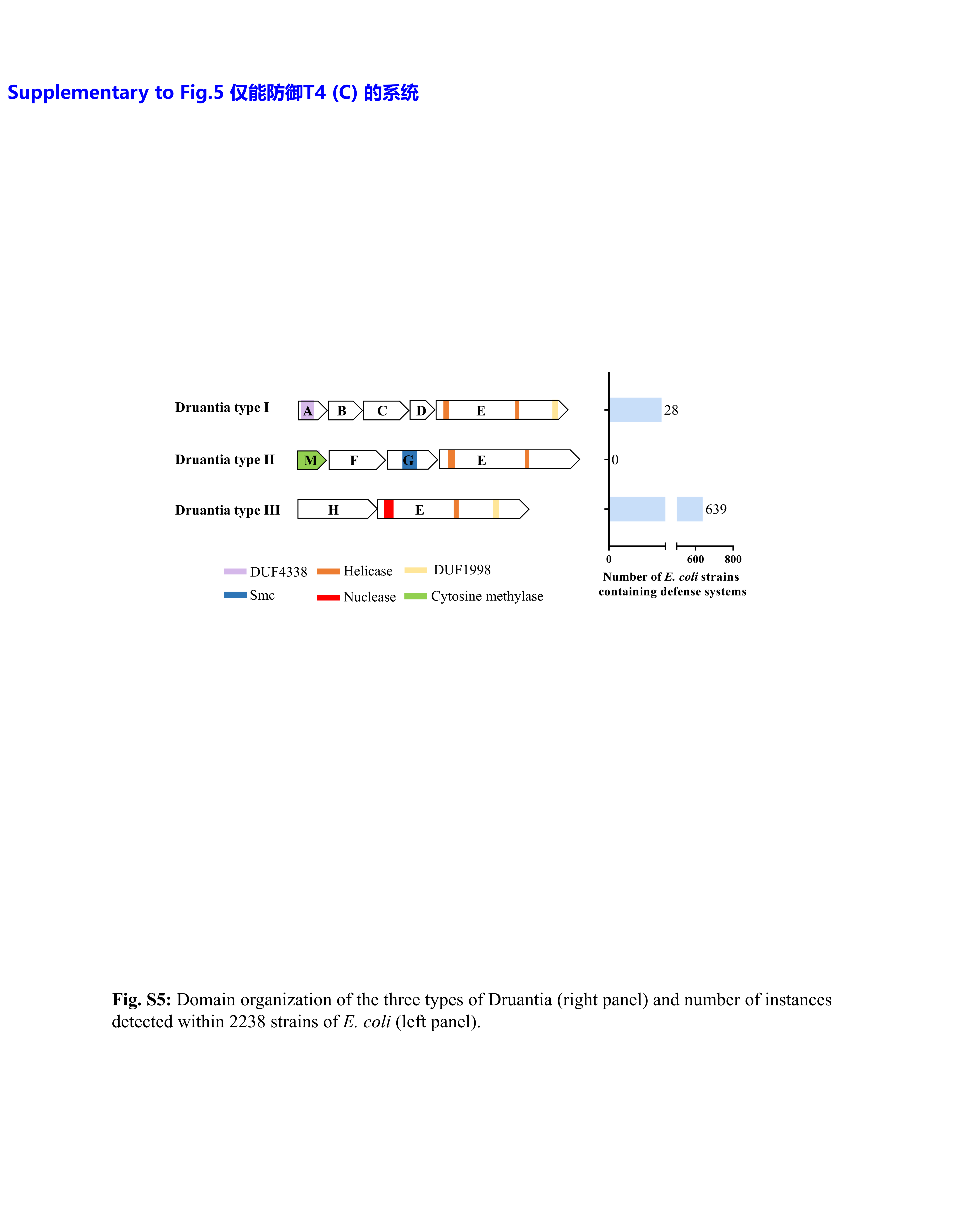
**Fig. S5**

**Fig. S5. The abundance of Druantia defense systems in *E. coli***. Domain organization of the three types of Druantia (right panel) and number of strains containing Druantia system detected in 2238 *E. coli* genome (left panel).

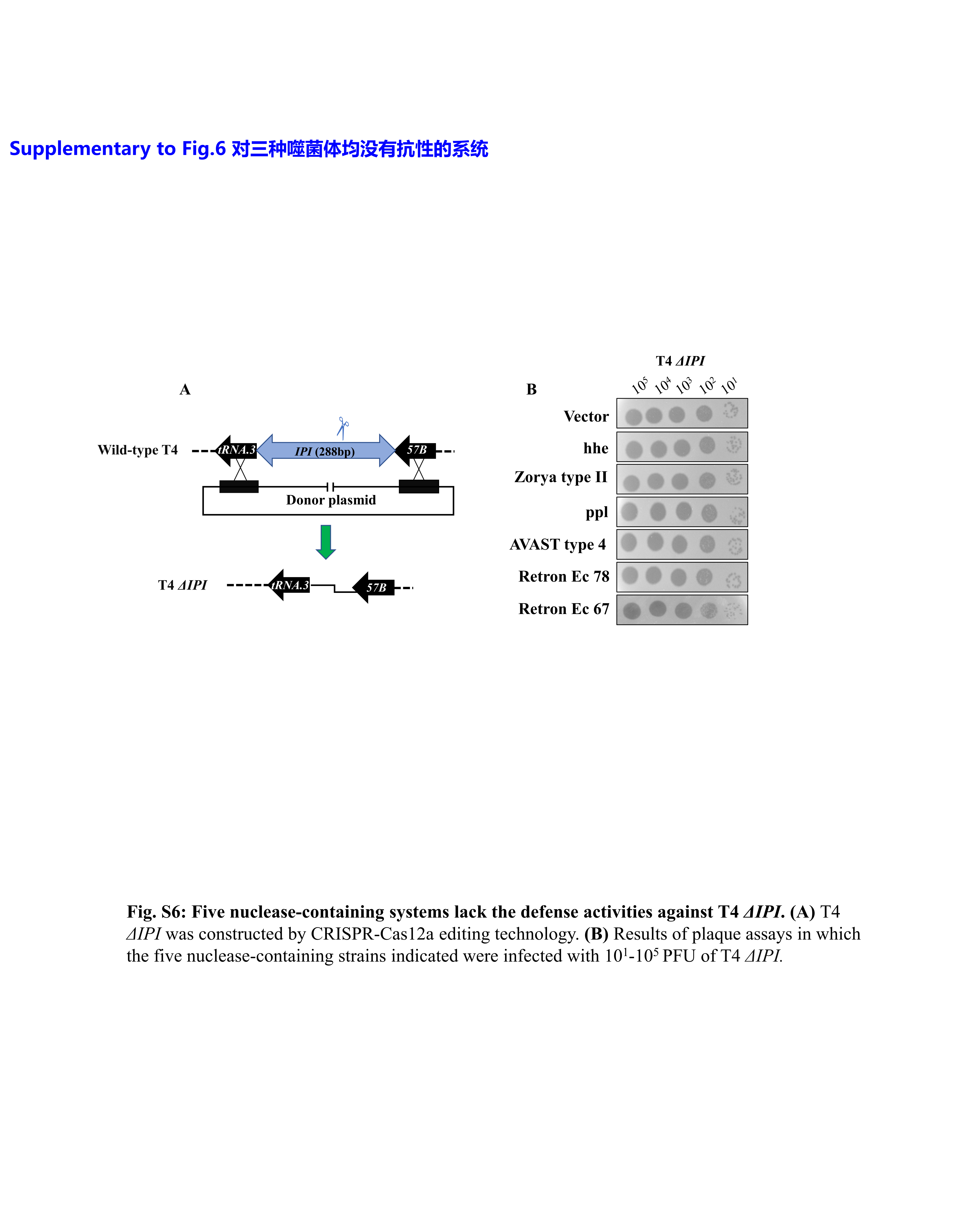
**Fig. S6**

**Fig. S6. Six nuclease-containing systems lack the defense activities against T4 *ΔIPI*.** **(A)** T4 *ΔIPI* was constructed using CRISPR-Cas12a editing technology. **(B)** Representative results of plaque assays. *E. coli* cells expressing the nuclease-containing defense system were infected with 10^1^-10^5^ PFU of T4 *ΔIPI* as indicated on top*.*

**Table S1.** Nuclease domain-containing defense systems reported in the literature

| **#** | **System** | **Domain annotations** | **Source** |
| --- | --- | --- | --- |
| 1 | Druantia type III | DruHE | *Enterobacter radicincitans* |
| 2 | Zorya | ZorABCD (type I), ZorABE (type II) | *E. coli* |
| 3 | Septu | PtuAB | *B. subtilis* |
| 4 | Gabija | GajAB | *B. subtilis* |
| 5 | AVAST type4 | Mrr-stand | *E. coli* |
| 6 | qatABCD | ATPase+QueC+TatD | *E. coli* |
| 7 | hhe | DUF4011-helicase-Vsr | *E. coli* |
| 8 | Retron Ec67 | Retron-predicted endonuclease | *E. coli* |
| 9 | ppl | PHP-ATPase | *E. coli* |
| 10 | mzaABCDE | MutL+Z1+DUF+AIPR | *S. enterica* |
| 11 | - | Restriction-like | *E. coli* |
| 12 | Shedu | SduA | *B. subtilis* |
| 13 | Ssp | SspABCDE, SspBCDE, SspABCD-SspFGH | *V. cyclitrophicus, E. coli* |
| 14 | Retron Ec78 | Retron+ATPase+HNH | *E. coli* |
| 15 | Ago | PIWI | *Thermus thermophilus, Rhodobacter sphaeroides* |
| 16 | Retron-Eco8 | Endonuclease+Retron | *E. coli* |
| 17 | - | DUF4297+HerA | *E. coli* |
| 18 | upx | DUF1887 | *E. coli* |
| 19 | AVAST type3 | DUF4297-stand | *S. enterica* |
| 20 | DISARM | drmA, drmB, drmC, drmD, drmE, drmMI, drmMII | *B. subtilis* |

**Table S2.** Sequence alignment of *E. coli* defense systems Gabija, Septu, Shedu, mzaABCD, and Druantia type III with corresponding systems reported in other bacteria

| **System** | **Reported from**  **source** | **In this study source** | **Query cover** | **Identity** |
| --- | --- | --- | --- | --- |
| Gabija | *B. cereus* HuB5-5 (AHEG01000016) | *E. coli* (EPEC47) | 23% | 32.86% |
|  | *B. cereus* VD045 (AHET01000033) |  | 36% | 28.57% |
| Septu | *B. thuringiensis* HD12 (Ga0125139_15) | *E. coli* (EPEC15) | 56% | 22.75% |
|  | *B. weihenstephanensis* KBAB4 (NC_010184) |  | 12% | 47.62% |
| Shedu | *B. cereus* B4264 (NC_011725) | *E. coli* (41-1) | 0% | 0% |
| mzaABCDE | *S. enterica* (NCTC5773) | *E. coli* (48-1) | 99% | 93% |
| Druantia type III | *Enterobacter radicincitans* DSM 16656 (AKYD01000027) | *E. coli* (EPEC42) | 83% | 32.65% |

**Table S3.** Sequences of *E. coli* defense systems Gabija, Septu, Shedu, mzaABCD, and Druantia type III

| **Systems** | **Sequence (5’ to 3’)** |
| --- | --- |
| Gabija | CTATGTCTGCATGAACATCGTTACTGTCAGTAAGTATCAAGATCATTGTTAGGATGTATCTGAATTTTTTTGTTTACTTAATATCCGATAGTTATGTTTCATCGTCAATATTAATTTTTATTGAATATAAGGTATTGATTTAAAAGTAGTTTAATCACCGCATTGGTTTTCATAATGTAATTTACATATGGTATTATATCATTTCTTTTAGTCTTTCTTGTAAATTACCTTGCTTTTACTTGACTTAGTCAGACTTTTGTACTCGATAGCTTTGTGCTTCTTGTGGCTATGGAGTAATCTAATGTGTAATAAAACTATGCTAAGGAACCAATCGATTACTATGTATATCAATAAATTAGCTATTAGAAATTATAAAAACTTTCGAAGCTCAAATTTCTGCTTTGTTAAAAACTCAGTGAATACAATTATCGGTGAGAATGCTTCAGGTAAAACAAATTTGTTTAATGCTATGCGATTGATTCTTGATGACTCACTTCCAATGAATTCAAGAGTTCTATCAGGTGAAGATTTTTATCGTGGCTTAAATGACCCCTTCGGGCACTGGATTATTATCACAATGTATTTTGATGATTTAAGTGAATCTGAAGAAGAACAGGTTATTGCGAACTATACAATAAATAATGGTAACGAAGAGGCATTAAAAGAAGGAAATTATACATTTATATATCGCCCAAGATTTCATATTAGGCAAAAGCTATACGAGTTAACTGTAGATAATAAAGATTTAGAATCAAGATTAGAGGCTTTTACAGAGTTGAAATCAACTTGTATAATATCCAAAGAGACTTACGAGGCTGTCGCTTTTGTTAGAACAAAAGTTGATTTTAATAATGACGAAGTTTATAAGCAAGTAGTTGGGGATTTTGAAAATTACATTTTTTCAAATCCCAATGAAGATGATGCTACAGTCATTGGTGTAAAAAAACCGCCATACTTCGCCTTAGGTAGTGAAGTTGCTTGTACTTATGTCAAGGCTCTTCGCAACGTGGTAGCTGATTTAAAGTATTATAAAACTAATCCACTGTATAAATTGCTCACTTTAAAAAGCAAACAAATTGATGATAGAAAAGATATTTCTGAGAATGTTAAGAACATTAATGAACAAATTTCAGCAATTCCTGAAATTGCTCGACTAAGTAATAAAATATCAACATCTTTACTTAATACGGTTGGCTCAACATATTCGCCAAAAATATTGGTTTCATCACAACTACCCGAAGATTTTACTGAATTAATCCAATCACTTGGATTAGTAGTCGAGGATTCTTTAGATTATGACGGTTCAGGAAGAATTGATGATTTAAGTCTTGGGGGGGCAAATTTAATATATTTGGCTTTAAAATTATATGAGTATGAAGAAATTAGAGATGCTGAAGAGCATATAACTCATTTCCTATTGATTGAAGAACCAGAGGCTCATATACATAATCATATCCAGAAAACTTTATTTGACAATTTTAATTTTCAAAATACTCAAGTCTTCGTTTCGACTCATTCAACCCAAATAAGCTCTGTTTCTAAAATCTCGTCGATGAATATTTTGGCTCGGCAACGAGGATATACAGATATCTATCAGCCTTCCAACGGACTACAGCCAAAAGAAATTTCGTCAATTGAAAGATATCTTGATGCAATCCGTAGTGATGTGTTGTTTGCTAAAAGTGTAATTCTCGTAGAGGGTGATGCTGAATTAATAATAATCCCAGAACTTATAAAGGTCACTTTAGGTATTAGCTTGGATGAATTAGGTATTAGCTTAATAAAATTGGATGGGACTGTATTTAAGCATATTTCAAATCTTTTTCATAGTAATAGATTAAAACGCTATTGCGCAATTTTAACTGATAAAGATTTGGCATATGTGCAAGAACCGGATGAACTATTTGATGCTGATTTCGTTGAATCTTTGAAAAACGCTGAGACTGACGGTTTACGCAGAGAACGAGAGTTGGCAGAGTATGTAGAAGGCAATCAATTTGTCTCCGTTTTTTATGCAGAGAATACGTTTGAAACAGAGCTTGTTCGGTATGATGAAAATTCCGATCTTTTCAACAGTGTCATTAGGACAACATATAAAAGATCAGGTGATATTGAGCGAGTTGTTGCAGGTATTAATTCCGATGATTTACGGGTAAGATTTAGAACTGCTCTTTTTTTCGCCAATAAAATAGGTAAAGGGTGGTTAGCTACTGAAATGGTAGAGTATCTCCATAATGAGAATCGTGTTCCTGACTATATATTAAAAGCGATTCATTTTGCACTTAAAGATAGAGAACTGAATGCTGTGCTTACTAAAATGCTTGCCTATAATATGTCTTTAATGGGAACTCACTGGGATGATGTATGCAACAAGATTAATTCTGAACCTGATTTTGATAAAAAGATGCAAGAGTATAGGAAACATTTCAATGATAGCTTTTCTAGATTTTGGGAGTTGTAATTATGCCATATAGAAATTTGACATCTGAACAACTAGATGCTGTAAAATTTGATGGCAATCTATTACTTACAGCTTGTCCAGGTTCAGGAAAAACAAAAACATTGGTTTCCAAACTTTGTTTTATCCTTGAAAATAAGGAGTTGTTTGATATAGGTAAGAAAAAAATTGTGGCATTGACATACACTAATGTTGCAGCAGATACTATTCTCGAAAGAATTATTTCATATGGTATAAATAGCGATTCATTGTGGATTGGAACAATACATAGTTTTTGTTTAAATTGGATTATAAAGCCTAACATTGATAGAATTCCAAGGCTCAGCAAAGGTTTTAATGTAATTGATGAGTATGAAAAAGAATTTATATTAAATGATCTGAGAGAACGTTATAACTTATCTCCTTACGATAAAATAGATACGCTTTTAGACTTAAGTTTTACTCCGGTATATTCTCGAAATACTGCTGAATATTCAGTGGTAAAAGCCTATCACTCCTATTTGTATGAGAATCGATTAATCGATTTTGATTTGATTTTGAATTTAACGTGCAATTTATTATTGAAGAATGATGTGCTATGTGAAAGACTATCTTTAATTATGCGATATATTTTAGTCGATGAATATCAAGATACCTCTCATATTCAATATGAAATACTAAGGCTGATAGTAAGTAAAAGAAATTCCTTGGTTACTTTTATTGGTGATAAAGAACAAGCAATATATACTGGATTGGGAGCGGTTGTTAAAAATAGAGAAGAGTTGTTCGATTACTTCAACCTTGAAGAATTAGAAGAAATGAGGTTAACCGGTTGTTTCCGTAGTAGTCAATCGATCATAGATTTTTATAGCAAATATAAAGATGATGCCTATTATATTAATTCATTGTCAGAACTGAAGACATTTCCTTCTGTGATACTCAAGGAAAATAGTGTTGATATCACCCAGTTGTCTGTTTATGTTTCAGGAATAATAAGAACTCACTTAGAACAAGGTATTTCTCCTGGTGAAATTGCTGTTCTATGCCCAAGTTGGTTTGATGTAATAAAATTATCCAATGAAATTATATCACTAAATTCTGATCTGGATATCGATGGTATCATGGTTTCTCCCATACCTAAAAATAATGAAAACTTGTGGCTCAATCTTGTAAAGCTTGCTTTAATTGAAAAGAATCCATCTAATTATCTTGTTAGGCAAAAACTTTTGCGAGATTTACTCAAAGAATTAAATGATTTAGAGTCATATACAAAATTCTTATCTCCAAAGCAAATCATAAAAGAAATAAATAAAATAGAATTTGCATTGGATTATAATTGTGAAATAGATATTTGGATAAAAAATCTTATAACTAACTTCTGTAGTTCATTAGGATTGGATGTTAATAATCAAAGCTATGCTTATAAAGAAATGATCTGCTTGATAGATGCTACTAAAAAACGAATGGCTAAATATAGTATGAACTATAGGGCAAATGATCTTCATTCTTTTTTTAATTTTAAAAGCGGAGTTAAAATTACGACTTGTCATTCGACAAAAGGAGATGAGTACGATGTCATTATTTGCACCGGGCTATTAAATGGAAAAATACCAAATTGGAATGACATTATAGATTGTAGCCCTGCACATCAAAATTATGTGGCTCGCAGGCTCTTATATGTCATAGGGTCTCGTGCAAAAAAACATTTATATTTGATATCTGAAAGCGGTCATAAGACACAAAAAGGTTATCCATATCAGGTCACTCCACAGCTTTAAGTACCCTTTGGTAGCAATAACTTTGGTATTATTGCTACGGGGAAATATAGTACTTATTTTCTTTTCTGCTTTGTCAATTGTTTAGGCAGCACAAGCTTTTAACCTTTGAGCTTGCACATAGCCGCTCCACCATTGCATGAGAACCACTCGTTCTGCGAGATAATCTGCGCGATTATATGCGGCAATTATTTCATCTTTCTTCGAGTGAGCAAGGGCCGCTTCTAAGACATCTGTCCTGAACTTACCCGACTCCTCTGCGGCCGTTCGTGCAATAGATCTCATTCCATGGGCTACAAGCTCTCCACCAAAGCCCATACGGATTATGGCCGCATTCGCTGTTTGTTCATGCATATGATTGAGTGGAGCTTTGATACTGGGGAAAACCCACTCTCTGTGCCCACTGATGGCTTTCATTGAATCCAATATTCGCAAAGC |
| Septu | GCAGTTCTTCGATGTTCATCATCAGAATCCTTCCGGATAATTAGCTCTCCCCTTTAAGGGACCATCCCTCTTATCCCTGCGCGCTACTTAAGTATTTTTGATTCTATTCCGGCGCCGTCCAGAACTTCAAACGCGTTGAAAATAAAAACAAAAACCCGCCGAAGCGGGTTAAGTGCGGGTGCGTTGAGGATGCCTGCCACATCAGAGGTGGCGAGGGATTTCTCCCTCGCCGGGTCTCTTACTCCTCAGGTTCGTAAGCTGTGAAGACAGCGACCTCCGTCTGGCCGGTTCGGATTCGTACCTCGCAGAGGTCTTTCCTCGTTACCAGTGTCGTCACTATGACGGTTAAACAGATGACGATCAGGGCGATTAACATCGCCTTTTGCTGCTTCATAGCCTGCTTCTCCTTGCCTTCCGGCACGTAAGAGGCTAACCTACATGTGTCTAGCATGAAATTGGCCTCAGATTAATGTTAAGCGTCTTGCAGGACGCGTAATGTTAACTGGGGCTTTTCTCTATCTGCCTTTGGTGTTCATGCCTGAGACAGATAGCCTCAAGCACCCGCAGCCATTCTACTTAACTCCCGTTACCCCGCCAATATGAAATCAGTCAGAAAGGCGATCCATAAGAATAATGAATTCTGTAAGGCAATAGTCTCACTGTCTAACATATCTTACCTGATTACTTTACAATATAATAATTCATAGTTAAATCAAATGAATGTCATCATCCAATAATTCATTAGTAGTAATACTACCTTTTTTGAAAAAATTTTCATTTGCCTCAGCAATTTTTGCATCATTAACAAATTCATCACACATCCCAAAATAATGAAAAAATCTATTTAGTTTACATATTCCAATGGTATAGTGCCCTTTAGGTGTTTTGTCCACATAAACACGATTTGCTTTTATAATATGCATCTCATAATCATCTATATGTGGATGAACGATGCGAAATGATGCAGATGCAGTTGGATAGTTTTTGCGCCGCCCCCCATTTATTACGGGTTCAAATAACACTTCATTTTTACCTTTGAATTCATTGCAGTCAGGACATACTACACATAGATTCTTAGGTTCAAAAATAAAACATAGATATTGAGATTTTGGCACAATATGCTCTATAGGAGCAGCTTGTGCAGCTCTTACCCCAATAGGCTCATGACAATATACACATTCTAGCTTTTGTTCTTCTCTGTAATGATTTCTAATTTCTGCACGGAGAACTTGCAAATCATCATCTCCCCAGTTGGTATGCTTAAAATCTGGAGAAGACAATTTTTTCTCAATAATATCTAATGACTCTGCGCTGAATCTAATTGGATTTTTAATTTTTGGCATAAATTTCACACACCTGAGATACGGAATTTACGAGTTCATTAATAGGGTCATTACTATTAACGTTTTCAAGGAGAAGCTGGAGTTGTTTTAATTCACGCCAATTATCATCTGTTACTAACTTAGCAGTCTTAACTTTTGAAAGTAAATTGAAAGCTAATCTGGCAATGTATTCATTCATAATGCCAGGGGCATCAAATAGTTCCGCAAGTTGATAGTCTGCAGACCGGTTGCTAAAGTATTTTGCTTTATAAAGATAACTTTTCGACAAAGATGTTATATAACAACCTTTATTCGGCAAATTGGATATTATTTGTGGTGAATGAGTTGCAATTATGAATTGACAACCTGAGTAAGTTAAAAATGCTTTTGTTAACATAATGATAAACTCTTCCTGCCATCGAGGATGCAAGCTTATTTCCGGTTCATCAATTAGTATTATAGAACCATCAGTTATATGCCCTGCAATTCCAAGCAACATCACTAATAAACATTGTTCTCCCGAGCTCGCACGTTTAAGTGACATGGGACCATAGCTTTTTTTTATGAGTCGCATATCCATCAATCTCATGAAGCCAGCGTTCATAAGTTTAAGAATTGATCTAGAAAAATAATTATTCGCATAAGTCGAGCCAACCAATGTTTCACCATTGGTAAAATCGACTGACAAGGTTACAGCCCTACGTTCAGGTTCAAATCGATTAATCTCAACCAGTGCTTCGTAAATATCATGACGTGATTCGGAGTTTAACTTTAAGAGACTCTCAATATTCTTCCTGTCAATTTTTATATTGTATTTTAAATATAACTCCTTAACTTCCGTTAATATCTCATCTGAATATACACCAATGAAGTTATAGTCAGGAGTGTCCGGTCCGTTGTAAGTTGGTTTAAATACAAACTCTATATGGGGCTCAAAATCAATAGAATGAAAAACATCGAGTAAATTTGCAGGATTTTCTGAATAAGCAAGTTTCTCTAGCCAACCTCTCGCTGCTGAAGATAAAAGGGATACGGCATTTGATACGCCGAAGGGGCCTTCTCCACGCATACCCACATAGCGATAATTATTTTCTTTCCTTTGGTGATTTAATGACCTACGCCCTGGAGGGAATTTATCGAATGGGCTGGTAGATACGGCAATTAACTTTGGTGTTATTTCTCTGTCAAATAATGGATTAAAAGTTTTCAAGAAATGAATATCTTGAATACAGTCTTTAGCAATCTGAGATAACAATCTGCTTTTTCCAACGCCATTTTTACCTACAATTACTGTAAAAATATTCGCATTATCAGAAACATCACCTTCTATAGCAAGTTTTAAGGTTCGACTATCATGATGGGTATAATATAGTCTTTTCATTTTATTTTTCTCTCATTAAAATGCTCTTAATTAAGAATACCTATGCCTTTAACAGGCTCATAGTTATGATTTGTGATGCTTTCAAAAGGGTATGTAACTAAGTATTGAACTCTACACCTAAGATTCTAAGATTTTTAAATTAAAGTAGTTAAACATGCTTTTGATTGTACTCATTCTATGTCCATCGTTGCTGCATGTCTAGATATCGTGCCCCGCTATTGTTATGTAATGATATGATGGACGCCCTGTCTGAAGCAGCCAGTGAAATATTTTTACTGTCTGCGTTTTCTACAGGAGTAAATCACATGAGCACAATCAGAGTTGTTGGTATTGATATTGACCTGTGCCCAATTTTTTGAGACATATATTGGTACGCTCTTGGCACAGGGCTGCTGCTGCAGCCCTGACAGCTTTAGACTTCGATCGATTCAGAGATCTCTAATGCATCATCAACTTCTATACCCAGATAACGAACTGTGCTTTCCGTTTTCTTATGGCCCAACGGAAGTTGGATCACCCGGAGATTCTTAGTTTTCTTGCAGACAAGGCAAGGTTTTGTTCTTCTCATGGAATGTGTGCTGTAAAGCGAATCTTCGAGACCAAGCTTTTCTCCCCCCTATGAAAGATTCGGTTATATTGCAGGATTGATATGTGTTGGTTAGTACCGACCCGAGATCGGAACAAGTAGTCTTTACTGCGTAAATTGCCAAGCTTTATCAATGCAGAAACAGCTTCTCTTGTCCCTTTGGTTATCTCAAATTGGATAGGACTGCCAGTTTTTCAGTTGCAACACCGTAGCTCTGCTTGAAACCGAGCTAAATCACGCGTTTTACCTTCCAGTTCAAGCCGGATTTGGAGCCCCCAGATATGAGATATCTGAAGTGGTCTTTTTGGGCCTATGATACTGTCTTTGTTCCTCGGTGACGTATTCATACTCAAATCTCCTGCAATGTGGAAGATTTGAGTATGGTTGTCTCTCAAGAGCGAAATAGGTCAACTATAACTAGTTGTTGTGCATGTACGAATTCTGCCAATTAATTATTGACAATAAAAAATTCTGGAATGTTCGAAATCATCATATC |
| Shedu | CTTCAAGGAAGCTCTGAGGCCATCTTTGGTAAAAGGTGAACTTGTCGGAAACCTCGCACTCACTGACCAGCAGCGCTGTGTTGCAGTTAATGCAATCAAAGCCTGAAAAAAATTCTGAGAAATGCAAATAACTTCAATGATATTATATACAAAGAGAAACGTCGGGTAGCCGGGGTAAAGTCCGGGTGTCACAGAACCCCGAACCGCTTCGCGGTGACTTCTTACCACAAACCAAGTCTATGTAAAATTTATTGCGTAGATATGAATCTAATCAGAGCCTTAAATTTAAAGATGGTAAATTATGTATATGACTGGTCAAGAAATAAACGATAAAAAGAAAGAGTATCTTGAATTACTAGATCGAGAAGAGAATGAACAGATTTATCAAACATACCTTGAAGAAAATACAATGTTCATACCAAGAGAATTCGAGCAAAACCATGGAATTCACTTCAGTACGGTATTTAGAAAGTTGCCTCTATCAAGCGATTATAAACCTGATTTCGTATATTTATCAAAAAGCTCGGATAACTGGAATGTAGTATTAGTTGAAATTGAAAAACCATCGTCGAAATATTTTAAAAACAACTCTACAACTTTTCATGCAGACTTCAATCTTGCTTTGCAGCAAATGAACACATGGAGGGCTTGGTTTGATGATGAATCAAACAGAAATCACTTTAAAAACAATATATTGCAGGGATTTATAGAACCGGCACATATGGGGAGAAATCCATTTAATTTCAAGTATGTCCTTGTTCATGGTCGTAGATCAGAATATGAAAATAATACACAGAAAACTGCACTAATCAGAGGGCAACAGCGAAGTGACTTCTCAATAATTTCATTTGATAGTTTGGCTGAAAATATAGAGAAAAAATATAAGTTATATGTTGGAGTGAAAAAAAATAGTCACTATGAATTAATATCAAAAGAATTTGTTGATGAAGGGATATTTTCATGGATGAACATAGATTTTTTAGCAATTACACAAAGTATATATGATGATGCAATCGCAAAATCAGATAGTTGGCATCACTATATAATTAAAGAAAATGGAACCGTCAAAACAATGGATTATGCTCTCCCTCGAATTAAAATAATTTGAGAGTAGTAAAAAAAATAGGTAATGACTCCAACTTATTGATAGTGTTTTATGTTCAGATAATGCCCGATGACTTTGTCATGCAGCTCCACCGATTTTGAGAACGACAGCGACTTC |
| mzaABCDE | TTGGTCAGAATATCGTCCAGCCATTCCAACTCACTTTCGTTTAACGGTCCCGTTTTCATACGCTTTTCCTTGTGGATCTCAACTCGCCAGCACCTATCTTACATGCCGGTCCGTATCAGAGATACTTTTTGAGTGGCTTTGCTGGTGATTAAAAATTAAGGAGGGTGTAACGACAAGTTGCAGGCACAAAAAAACCACCCGAAGGTGGTTTCACGACACTGCTTATTGCTTTGATTTTATTCTTATCTTTCCCATGGTACCCGGAGCGGGACTTGAACCCGCACAGCGCGAACGCCGAGGGATTTTAAATTCTACGCAGGATCAATGAGTTACATAGCAGTGCATTGCGAAACTTTGCTGTAGTTGTCCTTTTACTTTACTGTAATTTGCATTAAATAGATAGGCATGGATAACGTTGAGCACAGAAAAAGCACAGGAGAACAGTTTCCTGCCTGAGAATATTTTAAAGAATGAGGATGTGAAGCGCCATGGGGCTACCGGCGTTGTATGGTGATGTATAATCTAAGTCTTCAACTAAGAAAAAGTTCGGTTTATTCAACAAAGCCCCGCTGCGAGCAATCCATAACACACACGCAGGGCTTCGATCATGAACTCAGTGGACAATTTAACTACTCATGTGATCTTCCATTTTTCCATGTGCGGAGAAATAGATTTTACTGCGGACATACTCCCAGCATTCAGGCTTCTTCGCCCACTCCGATATCATCCTGCCATTCGCGGATTCAGTCAGAAGACTATTCACCCGATGGGCTAAGCTGACAAGCCGGTTGCCCATCGTTTCCTCAATGCCTTGTTGATTCCAGATCTCTGCAAGATCCGTTTGGTTGCCATAAAGATGTGAATAGGCGGCCACAGTATAGGCGGCAATATTCGCCTTAAATGCGGGGAAAAGTGAATTAATCAACTTATAGGCTTTCTTATAAATAATGGCCTGCGCGATGTAATTTTTAAAACGCTCAATGTTCAGTTCATCCCCATAACGCTCAGTATCCTTGTCGATCATCGTCATTAATGCGGCAAAGTTCTTCTGTCCGCCGAGGCTGACTAAATCCGGACGCTGGAGCCAGGCACAATGATATTTTGCGAAATCTGTTTTCGTTATCCGGCGAGAGGCAGGAATCGCGTCTTTTAACCGTTTGATTCCAGCGGGTGTTTTACCTTCCCGTTCCAGCATAACCTTATAACTGCCGTTCGCTCGCTCGTAAAACCAACGGCTGTATCCGTCCGGGCAATACACCGTGTTTGCCATTTTTTCCAGCTGTACATGTACCGGTCGATTGGCTGACAGATCGGAAATATTGACTTTATTCTGGCTATTCGAGAAGCGCGAAATATCCGCAATTAGCATCTCTTCTTGTGCATTATTCGTCTGTTTCAGCACAATGACTTTTGCGGGTACACGCACGTTACGCAGATTGGTTGCCGGGAATTTCTTTTTGGTGAAAAACATAGAAGCCGTCGTCTGCCCACCGTTGACAATCTGCATCCCTTGCATCCACGCAATACCAGGACCGCCTCCCGGCGCTTCGCCAAGCCTGACCTGATCGGCAACAATCACAATGCCGTTGTTATAAGCCATAAAACGCTCAGGCTGCTCGCGTAAAGTGTCACGAATCCCTTTGTTGACTTTCCCCGTCTGGCTCAGAAATGAGCGAACGTTCGCTTCCAGAATCCGGTTGCCATATTTTTCATAGATAAATCGCAGCGTCTCGCCCGGAACCACCGTCAGCGCATAATCATATTCACCCATTTCATCCGGGATCCAGACACAGGGGAGTGCGCCCCCCGCCACCTCATCAAAATTAACCTGCAGTTCGTCGCGTGGCTTACCTTCCTGCCAGTGGTTAAACAGGCGGACAATGTCCATAACCTCTAATTTAATGGTTTTACCGGCCACATCGCGGGACTGATACCAGCGGGTTTTCACCTGACCATCGGTCAGAACATAAATTCTGATTTGCTCCAGTTCCGTGTAGGACTGTTCGATGGTCGTCACCAGTTGCCAGGCATCATTGGACTGATCGAGCGTAGATGATAATTTACCGTCAACGCACTTCTGCAAAAATTGAATGCAGTGACCAGCAATCGCTTTTGTCTCAGCATCCGGGACGTGACAAAGCTCGTCGCTACCGTGATAAATGCTGACAAAAAGGTCAAGTTGATCGCCATCTTCTGAGAAGGCATAACCGCTGAGACGCACATTGTGTCCACTGACTTTCGCCATAAAGTGACACGTCTCGGCATCATCGAATGTCATGCCGATATCGGCCATATGACGCATAACAATATCGGTAAAAATCAGTTCCGGAAAAGGCGACTCAACCCCAGGCTGAGCCATTTGATCCTGATATTCCTTGCGAATCTCATTCTGCGTTTGACGTAAAAAATCAATAAGTTCCATGCGGTAATCCTGCAAAGTCTGACAAGCAATGATTAAAATTTTGAGCCTGAGGAATAAGTGTCTGGAGTTCGAGTTGATAGCACGCACTCACCACCGGCAACGGGACATTGGCATGCGTCAAAGCGGGAAAATCCGCTTCAATCGGAAGTGCAAACGCCTCTGTCAGCAAGAAATGACGTGTATAGAATTCAGCATGTTCAGTGAAATATCCGACGTGACAAAGCGATCCCTCAAAAAGACAGGCCGCCGGAGCGTTTCCCTCAAACCGCTGACGAAGACGGTAAATGAGGTCATTCAGGGTCGCGCCGGTGGGATGCTCGCTGAAGCGCAAACCGCACAACATGAGCGAGCCAGGCCGCTGCCAGTCCAGTTGTTCAAGCGCGTGGATGGTTACGCAAAAGCCTTGCTCTGCTGTGGTGCTTTTTACCTCAATGGCGCGTTCATCGAGTTCAAAATCCTGTAGGCCATGCTGCGGTCCTACCCATGCTTCGACTGCCGTTGACGGCGCAAGACCGCTCTCGATTAACCGCTCCAGACACGTCAATTCCCCGATTACCCCCACTTCTTCTTCATGACTGAGTGGACGGGTATCGCGCTCCATAAACGCCTGCCATAAACGAATCCGACGGAGCAGGGTTTCAAACAGACGCGGTTCAGAAACGTTCAAGAGCGAATGGAGAATATCCAGAATCATCGTCGTAAAGACATCCGCATTCCCCGTTTGCTGACGACGGATCATCAAACCGTTCTGCCGCCCCTCCTCTGTTCCGGTCATCTCAACGCGAAATCCTTTACTGGACGGCAGCGCCGTCGACCCCGGCGAAAGAGGATGAGGAAACACGGCAATCAGCATTTCTTCATTATCAGGCTGACGACGCCCGGCATGAAAGCTACAGCCCTGGTGCGTAAAAAGCCGAATCCCGCGCCAGCCTTGAGCGCCGGGCTGTTCGTCAATCTGAGATCGATCTAAGGCTTTCCAGGCCGCCAGCAGATCATCTTTACTCAGCCGCACCATACTCTTGCTCCCACAGTACGTTGTTAACTTTGTACTCCACCGTTACCCCACTGAGGCTTCCCGGAAAACTGATGCCAAAAGCCACCACCGGCAAGGCATCGTCCTGATATTCCGCTGCCGACAACGCCTGTTGCGGGTCCAGTAAGTAAATGAGCAATAGCCCTTTTTGACGCTGTGCCGGGATGTCGTTTATGCCAAATCCTTTGATCCGACGCAGTACTACGCCACCAGGAATGACGGGCTCCTCTCGCCCTTCATTGCGTCCGGGATCGGCATGAAAAATACGCTGCGTTTCGGCCAACGCGGCAAGCCAGGTCGATTCATCGCAGTCAATCCCTTCATCGCGTGGGGAAAGTAAACGGCCAATAGAATAGCGGTCAGTGACCCCCTCAGACGCTTTACGCATCATAAGCGGGACGGGGAAGCCACAGACAGAATGATGACGATCGACGCCTCCCCCTATCACCGCCACCGTCCATTGGGTGAGTTCATCAACACGATTCATCCGCGTAATGAAATCCGCCAGCAGTTTGCTGTTGGCTTTCCTGGCCTGAGCGTGGGTTTCATATTCCGTTAAGAAATCAATGATCAGCTCGGGAGAGACGTTTTGCCAAATGACCCCTTTCCATTTTTCAATCTTCTCTGCGCGCCGTCTCTCAGGGATCGGCGCGCCAGAACCGACACGCCCGGTCAGATGCTGGAAGGCCGCGTAGTTACGCTTGTGGTACTCCGGTTCTTTAAACAACGAAATCGTTTCGACCACCGTGCCACTGAAAGAGAGCCATAGTGAGCGCGCGCTACGCATTTTTAAGGGGGATGTCACCATTAACACGGGGTGTGATTTCACTTTTAGCCCGAAATCACGCGGGGTTCCGCCGCTGGCGACCATATTGTCAAACTCTTCCCGCAGCTCTTCTGAGGCATCCGCAATGTGCTCAAACCATTCAATCAGCTCATCGGTGGTATACAGCCGACATACATCCAGATATCCCTGTCGATAGCCAAACCAGCGCCCCATCTGCATTAACGTGTCATACATGCGGGAGGCGCGTAAGAAATAACTGGTGCATAATCCCTCAAGCGTTAGCCCTCGAGCCAGTTTGTCTCCGCCAATCGCAATGACTTTTAAGCCTGTCGCACTGTCCGAATAATCAAGCGCATCCTTCGCCGTTCCGTTTATCATTCGTACCGACACGTTTTCGATCACGGTATACAGCTTGTCGACGATCTCCCGCCACGCAAAGGCGTCGTCCGGAATCTGCTGCGGCATGACGTCGCGAATCGTCTGACCAGTCGGCAAAAAGTCGGCCTGCCAGAGTGATTCAAGCTGATGTAAAAAAGGCTCGTGTCCAATCCTTCGCGTCAGTCGCTGTCTCACGTCCTGAATGTAGGCGTCAATATGCTCATAAACGACCGATTGCACTTTATTGAAGCGGGTCACATGGACCAGCATCGAACTGTGTTTATCTCCCTGACCGCGTAATTCTCTGACACAGCACGCCAGCAAAAAGCTATCGATGGCATGTTTTAACGAGTCCGGGAAATGGGTTAGCGTATCGAGCGTGGGATAGTGCGAGCTCTTATGAGAAACCGGCATCCACCCTCTTTTTCCGTCATCGCTGCAATGATCGCTCACCCGCCTGATCAAAGGCAGTTCACCGGTCCGCCCTTCCGCGGCGGCCCGACCAAATACCCTCGCCGGGCCGATGTAATTAGAAGGTGCGCCGAGGTTAATGATAAAGGCGGAAGGGAACAAATCCGGGCCTTCATCACGCGTTTCATTGCTCTCGTGAATAAAAATATTGGCAAAGGGCGTGGCAGTATAGCCAACATACGCCTTACGGCTGAACTGCATCAACAGCTTACGAATCAGGCTATTGATTGCCGTTGGCTGATGTTCAGCATCCGGTTTTCCATCGTCATCGTAGACGATTTCCCCCGTATCAACTGAGGCATTATCCGCTTCATCATCAATCATCAGCAGCGGTAATTCCGAAACAAAACGCTTTCCGGTGACGGGGTCAACACTGGTGGCAACATGGTTCTCAATCCAGGTATGCAGGCGTCTCAGAATAGATTTATTTTTCTTTACCACGAACAGCCAGGGCCGTTGCTCAGGGCTGATTCCCAGATTCTTAGCCACTCCGGCACTGAAGTCGCCCTTTTCAGATCGGTTGGTGACGTAATTGGGACGAATGACGGGATCGCTATCAATATCGCCCACCCCAATGATGGTCACTTTTTCTCTGAGTGGACTCGTCTCGTAGCCAAGAAATCCTTCATCCAGACGCATTTGGGTCTGCGAGCGGAGGTTATTATGCAAACCCGCAAGCACAATGATTATCTTATATCCCGCATCCGCGGCTTTACAGATTAGACCGGTATAGTGGCTGGTTTTCCCCGACTGAACATGGCCGACCACCAGACCACGCCGGTCCCAGCGTCCTTCCCGCCCTGGCTGCTCAAGTAATCCCAGAACACGATCCGTTGAACGGTCAAGGGTATCCAGGACTCCCCAGGGCATCGTTTTACCAAGCCATTGGCGATATCGATGCCAGTAGCGCCACTCGCGTTTCGCATCTGCGGTCAGCCAAGGTTGATGTCCTTCATCATTACTCAGAGAAGTATCTTCGCCTATCCAGATGCTATAACGGCGGATCAACTCTTCGACCAGAATCTCCCGGTTATCAAGCCGGCTCCACTCCGGTTTCATCTGACAAACCATGTCAATATGTTGGTGCAGAATGCCAGCGGTGATCGTGCTTTTATCCTGGCCTTGCAGGAGCGTTTGCACAATCTGCAATACACTCAGTTGCGTGTCATCCAGGGGATTCAGACTCATAGCGATTTCTCTTGTTGATCGTCGGGGAGTTGTGCAATTAATTCGGGATAATTATCGAAAGGTTCCATATTTTGCAGATGCTGTTTCGCCAGCGCCGGTGACATCGCCTGCTGACCAACCAGGGTCTGGTACATGACCTGCAATACGGATAACACCTCTGCGGGCGACGCGGTTTCAAAGCCTGTCCGCGGCGTCTCTTTTGTCTCAGCCGTATCAAGCCAGATTTGCTGAACCGGAACGGTTTCCTCAATCAGTCTTAGCATGGCCTGAATTTGTGGGGATAACTCACCAGCCTGCGAAAGGACATTGCTGATGACAGGATGTTGTAACGAAATCTGGTAACGAACACCTGATGACGTCTTCTGCGCCTGCCAGAGATGAACGAGTTCCTCGCCGGGCTTGCGCTTATTCATTTTCCCGCGTTTGGCAAATGTCCTTACTGCACGATCGCGTGTTGATTGCGCCAGCTGAGTTAACCAGGGACGCAGCGAAACCGGCGGGCGCGCCATCGATTTACGAATATCAATCTTCCAGTCTATGTCGGCATCATTAGGGATATCCAGACGGATCCGCGCAAGGCGGTGGGTTTCATCTTTCGTCCAGGCCCGGGGATTTCCAAGTCCAAGCCAGTTGCCCGCCACCAGCAATCGCTCATTCCGGTATACATAAAATCCTTGCTGAGCCGTCCAGCCAGCCGGTCCTTGAGCCTGTTGATACTCCTGCGTCGTCAGGTGATCCTGATGCGGCAGAACATGACACTCCACCTTCACGGCAGGAGCGCCTGGCGCCATTGCCGAAGGCGAATGCCAGGGCTTGGAAGGATGCCCGCTGAGAAAGGGATCCCAGCCTTTAATTTTGCGACCATTGAGGGTGAGAGTGAGTCGGGGAGCGTTCCCCTCAAGGAATCGGTGAAATACCATCGCCAGATGTTGTTCAACGCCATCCATCAGATTGAGGAAATCTTTCTCACCGTAGCCGGGGGTGACAATTCGGTCTAAAACGTCCCACAGCACCACCGTACCGTGGGAGTCAGGTTCCGCATTTGCTAACGCCTCCTGGCTCCCTGGGTCAGCACCTTCAAGCAAATACCAACCGTCGTCCGTACTGGCGGCGAGAATGTCCAGATCCCACCGCAGGATGGTCGTTATCCCCTCTTTTTTAGAGAAGACCGTCAGGCGACGACATTGCGAAAAGGAGGCGGTTTTGAGTCCAAGACCGAAGCGGCCCAGATCGTGTCCTGGACGCTTTGTCAGCGGGTTTTTGACCCCCAGCCGCATCGCAACATCCAGTTCAGCGGCCGACATGCCGCAACCATTATCCCGAACCACAATATAGCAATCCGACTCACGCCAGTGAAAAGTCAGATCGACCTTACGGGCACCAGCACTGATGCTGTTGTCGATGATATCAGCCAGTGCGGTGGCGGTGTTGTAGCCGAGCCCTCGCAGAGCTTCGATCATAATGGCTGCATTCGGTGGCGCATGGCGCATGGTCGTCATTCTTCTTCTTCCTGCATCTCGCTCAACGCGGTTTCCCACTGTTCGACGAAGCCGCCTAATTTCGACGCCCGGCTGTGATGACGGAGCTTACGGAGTGCTTTGGCTTCTATCTGACGTATCCTTTCTCGCGTGACATCAAACTGTTTACCCACCTCTTCCAGCGTGAAATCTTGCGTCATACCGATGCCAAAACGCATTTTTATGATCTGTGTTTCGCGTGGCAACAGCGTATTCAGGACCGATGAAATATGTTCTCTTAGCTCCAGAGACTGAACGGGAGCACTGGTGAAATCCCTTCTGGTCAGCAGCGCGTCTCCCGACAATGCTGCTTCGCGGTCCTGAGCGTCTTCATGAAAATCACCAATCAGCACCGTCTGTTCTTCATATTTAGCCATCTGAAGAAGTTGACGCTCTGGCAGGTCGGTCATTTCCTGCAACCGTCTGAGGGTGGGCGTTATCCCTTGGCGATGCAGCAACTGATCCCGACTGCTCCGCCAGCGCCTGAAATGCTCATAGAAGTGAACGGGTAAACGAATGAGCTGAGCCTGATCGGCAATCGCGCGGCTGATCTTCTGGCGGATCCACCAGGAGGCATAGGTAGAGAATTTGAATCCACGCCGATAGTCGAATTTTTCAACGGCCTTGATCAATCCCAGATTACCTTCCTGGATCAGGTCCTCGATATCTAATCCTCGCCCGCAGTGTTTTTTCGCAATCGCGACCACAAGACGTAAATTGGCGTGGATAAACCCCTTCCATGCGTCTCTGTAACTACTGACATACAGACTGACATTATGGCATTCATCAGGGATGAGCGTTTGGGCGCGCTCGACTATCTGCTGCAGGTCACGGTACTCCGGGCGAGGGATATGATTATCCTGGTACTCTTCAGCTTTCCCACTTCTCAGCAGAATGAGCAGATCGTTAAAGGTGATGGCGTCATCGTCATCGGCTGCGGAAGAGATTTCCTCATCCGCTTCTTCCGCGTCGTCATCCTCTTCAGTAATCTCAGGCGATGTCGCGGGAAGGCAATTCGGATGCAGCAACTGTAGATATTTTGCTTCGGGTAATCTCGCCAGGATCGTCATCAAGTTGATAAGCGCCGTATCCATGCGCTGCCCGAGTCGCTCTTCCTTTGTTTTATCGACGACATCCCATTGGCGAATGTCCACAAGATAACGTTCAAAGATGCCGGACTGCCGCAATGCACCGAAAGCGTCAAACCAGGGCTCGAGAACATCACTCGCCGCTAATTCCCCAAAGGCATCACTTTCCGGGAAAGCATCATCGCCATCCACAATCAAAGGTAAGTCCTTGATCAATGCCTCTACGCTGAGCCGGAATTCAGGCCACGGCATGCCGAAATCATCTTCCCCGGCAGACCGAATGTCACGAACAGAGATACGACCGACATCCAGTGCGCTAGTGAGCAACGTTGAGAAATGGGGATAATTTTGCGGAGAGTTAGAAACTGGCGTCGGCGGGTCAGGAAGGGAGAGTTCTATATCGATCCACTCAGCAGCGTTATCTATCGGGCGATAGCGGGATAATAATTGCTGAGCCTCAGCGGAAGCCTGTCGTAATGTCAGATTATCTTCGGCGGGTTCCGATTCGATTTCAGCATCCCACTGCATAAAATCCATTGCGTCTTCGGTTTCGGGAGGTTCTGTTTCGACATCGGCTTCAATTACAAGCTCAACGTTTGCCTCATCATCTAGCCCCCCTGCTAGTTGCTGCCCGTCAGGCGGAGGAGATTGCCGATAACGAGCTAACAGATCACGCAGCGTTTGCTCCTGCGCGAGGTCACTGGCGCGAAACCCGAGGTTGTTTTGATGCTCAACATCGACACCCGCACACAACAACATTTCGCAAAGAGCGAAATGCCCATTCTGGGCCGCAATCATCAATGGGCTATTTCCCAGAGAATCTCGTGAATCTTTCGGCAATCCCTTGCTGAGATGAAACTCCACCACGGAGGCGACCCCGCTTTTTAGCGCCCGAATAAGTAAAGGGGGGATGCGATAAAACTTAGAAACAGAGCGATGCATGCCGACAGCCTTTCCTTGTAAACCACACCCGCGGAAATGCAATCTATCTTATAGGGAGAAAATCGAATTTGGTACTGTATATATAAGCAACATATCATTTTTGGTGATCTTCATCACCAATTGGGCTTTGATGAAAAAATCGAGCTTTTGCTGAGCTGAAGGACCTGTTAAAGCATTTGCTGACAACACATGCAGTCATGTAGTATACCAGAAGTTCAGAGCCACTAATTGGCACACTTACAAAAAGCCCTTCTGAACCTTGGCTCAATGGCTTTCTGGCTAGATGACGAGGCCATTAGGTCTCATATGAATGAAGTGGTTACTGAATTTGGCCACCTGAGCAGAAGTGCTTTACTCACCTCAGAACAACACAGGTAAGTCCCCCCTCTCACGGAGGCCTACCTGTCCTTTGCTATATTTATATGACTGCAAAGTTACTAGTTTGACTGGAAATCTTCGATTATCTTCTGATTGTAATCATCCAGATACTTTTGCAGTTCATCCCTATCGATCAGACGTACC |
| Druantia type III | CATTCGGCATCACGAATAACCACCCGCATACCAGGTGCATAATTTTGCTTATTCATCGGTTTTTCTGTCTATTTGGGGCAGTGGCGGTAACGTCATCCTGAAAAATGTGGCGCGGCACACTGCGATAAAAAAGAGGGCCTTAAAGATATCATGAAGGCCCCCGGCAAAGTGACAAAATGGGAGGGAATGCGCATAAATTGCGCATGATTTTCAAACAAAAATTATTAAAAAGTGAGCTGGGTCAACTCCCCTGGAAGATATCCCCAGGGGAGCAATTATTTATCCAGCGATAATATAACTCCTAAGGTCATTACCCGGGGCAACAGCTCGCGCTCGCGTGTTTGCCAGCCGTTCAACTTGCTGCGGTACTCGCAGGGCAAAATCAAAGAAGGTATCTTCAATACAGCGCCAGTAGCTAAGCCCCATGTCGAGCAGGACATAATGCTGACTGCCATCGCTGAAGGTGACTGTCAGTTCTCGTCCATGCTCCGTACTCTGGTTATCCCCCGTTAACCAGCTAATTGCTCCCGGCCACGCAGGCCCCAGATATCCTTCTTCAAGTAAAGTGGTCACAACCTGCTGACGATCTTCCTCATTAAGCCACGCGTTATAGAGAGCGTCGTGTTGGCGAGAGTTGTAATCCAATTTTTTGACGCAAACAGTGAGGCTGGCGCGTTCCGCCATTCCCTGTTCAACCAGCTCGGTTAAGATTTCACCTAACAAACGCGCCGTGAATGGGCTATTAAGATAACGATCGCTGTACTCAACATGCGTTATCTCTTTGTGGCGGGTAAATGCCTGCTTCCAGCCAGCGTGCTGTTGCGTGACTAACGACCAGAAACGCGAACCAAAACCTTCCAGCGGCCCGTCCAGTTGTTCACAGAGATTGATGCGCACCGCCCCGGTGGGTAAGGCGGGCAGTAAATCTTCTGCACTTAAGGTCTGACCCGAGAAGGTTTTTCCGGACGTTCCTTTAAGTGTCACTACTGGTGAAGACGCACTTTGTCCCCATAATTCACCAGGGGCAACGGTAGCGCTGTCGTCTGCGGCCCACTGTTGGCTTTGCCCTTCGCGAGTCACCTGCGCAAGCCAGCGCCCCTTTCCTTGCATGAGTTGCGCAGTGGCAATGGATTTCACTTGTAAGCGTCCTCCCGGCATCGCCGCCAACGCGGAAAGCTGATGTCTGGAGCTATCAGTTAAGTTTTCCAGCGGCGTGGCCACCAGCAACTCAACATTCCCACCGGAGGAGGCAAACTGGAGCAACTCTTTAAATAATGGCCATTGGGCAAAGTCCCACTGTTCCTGCGGCCCCGCAAGAACCAGCGAGCAGGAGTCATAAATCTCGCTGTTCAGGCATTGCGCAATACGTAAACTCAGAGGGTTAGTTTCAACCCGGCTACCATCGCCGAAGTACTGATATTCTGGCGCGAGGTTAAGTCGCTGCAACCGTACATTGGTAAGCAGGGTTAATGCATGGTGACGATCTAATCTGTTGACATAATGCTGACTGTCGTAGCTGAGCAAGCAGTGGTGACAAAATTTATCGCAACTGCAATCGAGCAATTTTCTGGCGTAATTAAAGACATCAGCCCAATGGTCGATCAATTGCACTGCATAACCTGCACCACCCGCATTATTGTCGTATATGAAGATGGAATATCCGGTATAGCCATTGATATCGCGCGCTTGCTGCGTCGTCACGCCCAGCTCAGTATTTTCGATACCCAATTTATGCGCTAACCCTTCACGCAAAGCGACCGCAAGGGAACGCGCGGCAACCTCATCACGGATCAACAGGCCATTATCATCATTAAGTTGCAGCTCAACCATATCTGTACGACTGCTATACCCCAACCACAGATCTTTATGCACATGACCGTGGCACAATTTATTGCTCCCCTGACGATCGTTACTGCCCCCTCGCAACAGGTAATGCGTATTTTCCCGCTCTGGGTTCTTTAGATTTTCAGGGGTACGCTGGGGTTGAGTTTGAGATTCTGCCCGACCACATGAGAGGCAAATTGCGTAGCCGTGCCCAAATTCCCCTTCACTGAAGTGGAACAATTCCCCACTATGGCTAAAACGGTAACGTCCTAGCCCAGCATCGGGTAAAGAAACCCACACCGAAGACGGTACAAGAACACGTGGATCTTTCCAGGGAAGTTGTGCAGGCATGGATACATCATTATGTGCCTGATAACGAATATCTGTCGCAAAACCAACAGGCTGTAAATAACGCTTCACATCAAGTTGCTGTATATCCGCTTTGCAGTGACTACATTCGACAGGATGGGTAGTACTGGTTGCCCCCGCACCACATTTACGGCAATGCCAGAACCACGGAATATGATGATCGCCTGACAGCTCCTGCCCGCTGGCAAGCACCTTGCCAAGCATAATGCCCGAGGAACGATATACTTTTCCGCCCAATACCACATCTGCACCAGGCGCATACTCGCGTAATGCTGTAGGCAAATCACGGCTGGGCAATTTTTCAATACGACGAGAAAAGCTTTCTCCCCCTTCATTAGGGTCAGCCTGAGTGGCTTTTATTGCGCGACGTTTTTCCAGCTCATCTGCCGTCAGATAATTGAAATTGACTACGCCAACCGGGAAACCATGCCCCGGCAGGAACGCTTCACTAATGAGTTTAGAAAACAGATATGCACCGCGATAATCACGAAGCTGATAAGCGATGGCTTTACCTGCTGGCGTTTCTTCCCACGCGCTGTTATCCGCCTTCAACGCTTCGATATTTTCCAACAGTATCGCAAGTTGGCTTCTCCAGCGGATATCAACCTGCTGCATCATTGCTTGTGTGCGGGTCAAGAGCTGGGTTGCCGTCAATGACATCAGGACGGTTCCCTGAGTAAGGCGTTTAAGTTTATCCGCCAGCTCATCGGCCTGGTGATTTAACCAATGAAGAAAACGAAGACACTGGCTCTCATCACCTTCAAAGAACCATTTGCAACTCAGTTTTGTCGCATCAGGAACTTCAGCTCGTAAAAACAGACCTAAAACCTGGGCATTAACATGTCGTTGAACAATGCTACTGCTCCCAAAACGAACCCGTGGCGCACTGGCAGTAGTGTTGAACGCCCAGAGGGGGTCTTTGAATACCTCCATTCCATGCGCCGTGTTTTTACATAACGTAATGGCAGCAGATGCACTCTCTCCACGACGCCCCGCGCGACCTGCACGTTGCAAATAATTTGCCGGCGCTGGAGGAACGTTGTTCATTGCCACCAAACTCATACCACCAATATCCACCCCCATCTCCATGGTCGTGGAACAGTTCAGCACATTAACTTTGCCTGACTTAAATTGTTTTTCATTAAACTGGTTCTGCTCAGGCGTTCTCTGCGCAGAGTGTTCCTCCAGACGATACCAGCTATCTTTCAGGGCCAGTCGGTCGCTGCGAGTAGACCATAATCCTTCCTCGCGGGCATGTTGAATCAGCGCATTACTTTCCAGCCACTCCAGACGTTCTTCTCGGGCGATTTCTCCACCACCAGACAAACGCCAGTGACGTACAGGAAGCTCAGGCATTTCAATCAACACTGCTTTTTCAGGTGCAATTTCTTTGCTACCCGGCAAATACGGAGAGTATCCCAGCAACGTGACGTCTAACGCACGTCGAGTATAGGGGCAACGCCACACTTGCTTTGGACTGTAAAATTCAGCTTGTTGTTTAATATCCAGCTGATAACCCGACTCAAACTGACGTAAGCAGGGTCGAATTGCATCCCAGACCTCAATCATTAAGTGGTTAACTAACGATTTCCAGTGAGCCTGCTCAGGTTGTATTGCCGGAAAGACTGCGATTAATAGCTTAATGACCCGGTTATAATGGCGATCGCGTATTTGTGGCCAAAGTTGATCACGTTTCTGCCCTTTGTTTTTTTTATCTGGCCCCTGGAGCATTTTTACCGGGAAACGTGCCCCCATCCAGCGAGGGTATTGCTCGTTTTCGTAAAAAACGGCACTGTTTTCACGGATAAAGAAATCAAGAGTAATCTTCAATAAATTACGCCATTCCGCTAATTGCTTATCGGGATCGGGTAATAATGTCTTCCAGTCCTGCGGGCAGGTTGTAACTTTATCAATGAATGGATAGTGAATGGCTACCAGACCCAGACTTTCGAGGGTCCAGGCATTTTTAGGGCGGCGACCAAACTCGCTGTATAAACAAAAATCTGCAAACTGTTGTTTAGTAAGATCAAGCCCAGACAATTCTTTAAAACTGCCTCTCATTCTATCGGCAAGGTCCACACTGCCAACCAGCTGTTCAGACAGCGTTTGCCATGACATGGTTTTCTTCTGCTGATTGCCAGAGTTAGCCTGAAGTATTTGCATCGCTTCAGCATGCTCCGTGCTTAAGGAACTCACCATCGGCTCAATAATTTTAAGTGCCTGAATGTCCGTCGGGCTAAGCTCCTCATTCGCAGCCATCGCTTTTTTCAGCGCGTCGATGGAATTTTTTAAAGGGAGGACAGAAAACTGGCTAATAACTTTCTCTAACAGTGCACAGTGTGCCGGGCTGAGTTTTGCTGAAGGCTCTGTAATATTTGCCAGTTCCTGGTAACAAAGAGCTCGCAACGTCGCTTTATCGGCATCCTGCTGTAAGCGGGCTGAAATGCGAGCGGTACCCTGACGCGAATCCGTAAAGGTGAGCAAGCGTCGACCATTAAACGGTCCTAACTGCTCTTCTCTTGTTTGCGGTAATGGAGAAAACTCCAGCAGAGTAGGAATAATTTCATTTAAAAAAAAGGGCGCGCCGAGCCGAAGCGGACGGAATAAAGGATTATTTTTCCGTTGAGTGTCACCACAGCAAACACAAGCAAAGGAGTTATTGCGTCCCTCACCTCGAGATTCCGGTCTTAACAGGTTTATTTCATAGTGCTTTTGCGCTTCACTATCAATTTTCCCCGCCGTCGTTTCTTCCAGTGAAATGATATATTTTTCACCATCGCTCGCAATCAGACGATCAAATTGAGAGTTATTGTCTAATTCGTCGTTGTCTTCATCTTCAGAATAAACATCTACATCAAGAGCAAACTCATCCACTTCGGCAGCGGCTGGTTGCGCTACCAACCAGTTCGTTCCATCACCGCGCATCTCCTCTTTGGCAGACAAATAAGCTGAACCACACCCGGAGCAACGTAAAACTTCAAATACCGGCGCTCCACAGTCACAATATTGGCGCTGCTCAAACCATACCTGACCAAATGGCCAGTCACTCTGATTTAATTCATGTGCTTTATGTGAGCATTGTTTGTTTGAACATGCCCATAGACCCGCCAGTGTTCTGATAAACCCATGCATACGTAATGGCGTAAAAGCCAGTTCACCTTCTCGCGATCTTGCCATTAGATCGAGGAGTTGCAGTAGCTCACGGTGATTAATATCTGGCCAGGTTCTTCTGGCAACATTGAGCAACTCACTCAAACGTACTGCCTGCCGTCCGGGATGAGTTAACGCCTGACGCAATTGCTGGGCCACACGGTAATGACAAAGCTGTTGGTAAAGATCCTGAGGGCTCAGGGATTTAAGAGAAGTTAAAGATGTGGGGTATTCATGACGAATCAGCGACTCCGATACCCGCGGTATTTGTCGATAACCGCGAACCACTTTGACATCACGTTCTGTAACGCCCGCGAGATCAGCGACAAATTTAGCAAGCTCCTTATTACCTTCAGGAGAGTCTTCACCAATAGTGGCAGAGGTCGCGATAATCTGCACAGGTTTATCTGTGCCGGGTCCAACATTGAAAGCCTGCATAACACGACGTAAAAGCAGCGCTAACTCAGCAGCCTGAGAACCAATATAACTATGGGCTTCATCCAGGACGATATACTTTAACATCCCCTGAGACTTTTCGATGATTGGGGCATCTTTACGACGAATCAGCATATATTCCAGCATGGTGGTATTCGTCACCATGATGGAAGGTGGTGCTTCATAAATGGCTTCCCGACTCAAAGCTTGTTCAGGTACTTTCAACTGATCTTCCTGAATTTCGTATTTGGTATGCCGGGTTTCGCCATTATATAGCGCGAAGCGTATTTTCCCCTTAAATCCGCGCGTCCATTCGCGAAGCCGCTCCTGTTGGCTGTTAATTAATGCATTTAACGGATACAAGAAAATAGCTTGTGTCCCTGTTAGTGACTGGCGGGTCGATTTTTGCTGACGAATAAGATCTTCCAGAACCGGTACCATAAAGCATTCTGTTTTCCCGGATCCCGTACCGCTGCTCACTAACAAGGATTTTTTGTCTTCCAGAATCGTTCGCCACGCAGTAAGTTGATGACGAAATGGCACTGGCGCTTTGCTCAATGCATCAACCAGCGGGCTGGAGATCAGCTTGTCCCGCAAATCCTCCATTGTTTCATCCGCAGGCATCCAGCCGAATGTACTTTCGAATACCGGGTCTGCAAGCAGTGACCCCTCCTGACCGGGAGGCTGAAAATATTGTGATTCCAGATAATGAGTCAACGACTTGCTCCGCAATGCCAACTGACTTACCACCGACCTCACCGCACGCTGATGAAGCTGATTTAGAAGTGGCGCATAATGCAGTTGCTGAACGTCCATCAATGACATACTCCTGCTTGTTCAAAATAATATTGATAGATTACCCCTAATTGCTGTGGGGCTTGCTGATAGTGAAATTCAAAAAGCGCAGACAGTTGATGTATCAGGTTATTATCTTTTTGGGTTAACAGCGCTGCGACTAACATCAAATTAAAACGTTTTTCAAGCCGTGAGTGATATTTATCTTCAATCCAAAGCTTATTAAAAACAGATTTTAACTCTGGAGTAATAGCACTAGACCAAAGTTCTTTAGGGAAGATACTGATACGCCCAAAGCTGTCGGGGTTCCTTAATAGCATCTGTTTAGCTTCATTAAACTGCTCACGACTAACAGTTGGCAATACATCACAACCGGTTGGTAGGCTCTCACTATATGTAAGTAACATATTCGCAATCGGAGCCAGCCGACGATCGATATGTGCACGATGGTTCATAAAAGCAATAAAGGGTTGCAATATATGTAATGCTAATGTTTTATCGTTAATCTGTTTCTCCAGATAAGTCCAACATTTTTGGAAAACATCTTTCCAGATTGAAACGGGAATCCATCCCCATGAAAAGGGTAATTCTTCCTGTATATCATAAATCGTATCGCTATCACCAGAAAGACATGATGATATTAACAATTGAACAACCACGCGGTAATTACTCGAAATACCTCTGAACAAGTCCAGATCCAGCAAGGATAAGCCATTATTCATTTTGTATTTCTTAATATACTCTGTTAATAAACAGCAATAATTTTGCAATGGGTTGTTATCTATTTCAAAAATTAGCTCATTAAAATTTTTTTTGCGCTGTTCTGGGTCTGCTTCGCATAGAGCATTTAGTAATAAATCATTTGTCTGCAATACAGGTAGCATCCATTTAATTACTGGTGCAATGCGAGCTGTACCATCAAGTTTTCCCACAATTAACCATGGACTATCCTGAACATTCAGAGCTGAAAGATCATAAGCAAACTCATCTTTTTTTAACAATGATATGTGTAAGCCAGGATTCTTAAGAGAAATAGCATCAACAACCAGTTCGTCTATTCTATTTGCAGGAAAAACAACAGAATTTTTTATATCAACCAGTAACCCTTGATCGTGTTCAATAAAACTACCATCGTAGCGTTTAATATTGAGTGTTGCTTTATCCTTTCCAAATCTGTCAGAGTAAAAATCAACACATAATGTACTATTGCTGTTCCATGCACATGCCAGCATAGCATTTAGCAAATTATAGTTCTCATAAAGTGAAATTTGTTGTAATTTGTCATTGCGTCCTGATAAAAGAGGAAGGTCGGCATGTAAATAGCGGAATTTTTCAGCTACTGGATTTTCATCCAATAACTCGATATTAAGATATCCTCTTCCAGCACATGATATGGTTAATAACTCAGCATCTATTCCATGCAAATGTGCTATAGATGCCACACCATCAAAATGAACAGAACCATCATTAGTTACAAATCGGCCACCGCTAACCGGCTTAGGTAATAATAGTTTAAGATTAGAACCATCTTTCCAGGAGACCCGTAAATCAACGGTATTTTCCAGCATCAGCGATGAATTATAATAAATAGCACACTCGTTTTGTGAATAATCGGACTTAAGTGTTACTTTTTCATTGTTTGAATATTTATCAATTCTAGTATCCGTTATACCAGAAAGCATAATAATGCCTTGCTGAGCACTCTCAGGTATAATATTAATATCGAAGTCTGCTGGTAAAACTACAACTTTACCGCTAAATAAAACCTCATCATTAACTATTCGCCGGACTTCTATCTGTCCTTTTGGCATTTCTGATGCGACAGAATACCAGGCATTATTACCTGAACGAATGGAACGCCAAAAAAGTTCTTTCTCCGGTACAATGCCATATTGCAGATCCTTTTTCCAACCGATCTTCGGCCAGGCTCGATGGACAGGATAATCCGATTTAACCAGTTCAACTTCTGTCGATTTAATATAATATTCAATCGCAGAATCATAAAGTTGCTGGGTGCGAATTGTACAAACAGCACCATCATGTAAAACAACGCTAAATACTCCACTTATTTTAGTTAAGCTGCGTTCACTGTTTTTCAGCAACCTTGGGATATCAAACTCACCTTCTCCACTAATATCTAAATGGCTATTTTTCGGTAGCGAAATAAACAGAGCATTTAGCCTGGAACTCACAGACCCCATACCAACAAGTTTTAGCTGCGATTCACTCTCATTCATCGCTTCAAAAACCCAGGGAAGTTCTTCAGTAAGTTCATAACCGCCTTTTGGAATTGTGTGACCTAACAGAATTGGACCTTCATGAAGCGACAATGAGATTTCGGCCATTGCATCTGCACCAGAGAGTTTTTGCATCGCAATAGGTAATAACTCGACCCGCCAATCTTGCTGCTCATAACGGCTTAGCATTGCCAACCTTGCCCCGCCATTTTTCCATTTTCCTGAAATGATGAGTCGGGTCTGCTCCGGCTGAATATGGCATTCAAACAAAGAGGTTAACTGTTCTGTACGCATTGTGGCCGGGAATCGGAACCGGGCATCGCAATACCAACTGTCATCAACATCGACCCAGATTCTTTCAACCTGAAGACTACTTGCACTGCGTATTTCAGAAGATTGAGAAAGTAATCGCCTTACAATCTCTGCAGCATTCGCTTCGGGGAGAACGAGTGGAAGTTGAAGATGCCAGTCTGGAGAGAGCTTACGTAATGCAGAAACGGGATCACTGCTATGAGGAGGATGCTCACTAAGTAATCGGCAGAGAGTCTCGCAGAATTCACCGGCTATTTCATAGACCAGTTCGTTTTGCATTGTGACAGGAATCGTATCGCCTAATTCTTGTGCAATTTTAGCGGCAGGATATTGAGATGATTGCCCACGAAGCGCCTGATATATCCTTCTGAAATAGGTAATCAAATAACCGCTCTCATTTTCAATCATGCGGATAGGCAAACCACCTTCACAAGCCAGTGTATGCAGATAACCCGAAGCTTGTCCCAGGTTTCTTATCTTTCTTTTCCAGTAGCGAATGCCATCATTAACCAGCTTCTGTCGATGAACATACGGAACTTTCCAGTTAATAGAATCTAAAATTGTGTCCCATTTAGGGTGTCCGACCGTATGAGTCCGGCGGATAAACTCTGCGGCATATATGCTAAATAGAACATCGCTATAGGAAATGAATATCGTTCTTGGATAACTACGGGGGGCATATGTACGCAGTAGTTCTGCCAAACTTTCGTACTCTGTATCAGTACAACGGTAAGCATATAACGCCCGTCCATCAGCTCCGGATAATGAACGGCGACTAAAAAACATGTTAAGCCATTGCAAAAGTGGGTTAACTGGAGTTTTCACAAGCAACCTGCCTTCCTTTGATACAATTCGTAACAGGTTACTATCATCATAAAAAAGCTCAACCCGATGAACTCGCTAAAAATGAGACAAATCATTTATATCTCGAAAAAACTTGTTACAATCATGAGCGCTACACCGAACTTAACCATATAAATTATGTGTGTTTTGTTTATTTTTTAAACGATTACAACTATCCATTATTTACACAGGTATCAAAATGTTAGCGCAGCTTTTTGAGCAGTTGTTTCAATCGATAGACTCTACACTGATCACCAATATTTTCATCTGGGCTGTTATATTCGTATTTTTATCAGCGTGGTGGTGTGACAAAAAAAATATACATAGTAAGTTTAGAGAATATGCTCCAACCTTAATGGGGGCATTAGGTATTCTGGGTACTTTCATTGGTATTATTATTGGTTTACTCAATTTTAATACCGAAAGTATTGATACCAGCATCCCCGTATTATTAGGTGGCCTAAAAACAGCATTCATTACAAGCATTGTAGG |

**Table S4. *E. coli*** strains containing 2 copies of the same defense system

| **#** | **NCBI accession number** | **Druantia** | **Gabija** | **Septu** |
| --- | --- | --- | --- | --- |
| 1 | CAADJH010000002.1 | 0 | 2 | 0 |
| 2 | AP022360.1 | 0 | 2 | 0 |
| 3 | NZ_CP026939.1 | 2 | 0 | 0 |
| 4 | NC_011601.1 | 0 | 0 | 2 |
| 5 | NZ_CP026755.1 | 0 | 0 | 2 |
| 6 | NZ_CP027587.1 | 0 | 0 | 2 |
| 7 | NZ_CP027597.1 | 0 | 0 | 2 |
| 8 | NZ_CP054236.1 | 0 | 0 | 2 |
| 9 | NZ_CP059840.1 | 0 | 0 | 2 |
| 10 | NZ_CP050201.1 | 0 | 0 | 2 |

**Table S5.** Nuclease domain-containing defense systems cloned in this study

| **#** | **System** | **Nuclease type** | **Source** | **Promoter** | **Genes** | **Length (bp)** |
| --- | --- | --- | --- | --- | --- | --- |
| 1 | hhe | Vsr: very short patch repair endonuclease | *E. coli* (EPEC40) | native | 1 | 6219 |
| 2 | Druantia type III | Type III restriction enzyme | *E. coli*  (EPEC42) | native | 2 | 10446 |
| 3 | Gabija | TOPRIM | *E. coli*  (EPEC47) | native | 2 | 4650 |
| 4 | Septu | HNH nuclease | *E. coli*  (EPEC15) | native | 2 | 3785 |
| 5 | Zorya type II | McrA: restriction endonuclease | *E. coli*  (EPEC41) | native | 3 | 4133 |
| 6 | SspBCDE | DUF1524: HNH nuclease, DUF262: ParB-like nuclease | *E. coli*  (22-1) | native | 4 | 9006 |
| 7 | AVAST type4 | Mrr-like nuclease | *E. coli*  (29-1) | native | 1 | 5797 |
| 8 | Shedu | DUF4263: PD-(D/E) XK nuclease | *E. coli*  (41-1) | native | 1 | 1237 |
| 9 | Retron Ec78 | HNH nuclease | *E. coli*  (42-1) | native | 3 | 4220 |
| 10 | qatABCD | TatD: magnesium dependent DNase activity | *E. coli*  (46-1) | native | 4 | 5635 |
| 11 | ppl | PHP: Polymerase and Histidinol Phosphatase | *E. coli*  (11-1) | native | 1 | 3392 |
| 12 | mzaABCDE | DUF4420: PD-(D/E) XK nuclease | *E. coli*  (48-1) | native | 5 | 10348 |
| 13 | Restriction-like | Type III restriction enzyme | *E. coli*  (EPEC23) | native | 4 | 11292 |
| 14 | Retron Ec67 | HNH nuclease | *E. coli*  (STEC138) | native | 1 | 2724 |

**Table S6.** Spacers used for construction of recombinant T4 phages

| **CRISPR-LbCas12a plasmids** | **Spacer sequences (5' to 3')** | **Targeted gene** | **EOP** | **Generated mutants** |
| --- | --- | --- | --- | --- |
| pLbCas12a-crRNA (*Beta-gt*) | agataccatgttcataggaattt | *Beta-gt* | 7.6×10^-5^ | T4 (α-ghmC) |
| pLbCas12a-crRNA (*Alfa-gt*) | ttacaccacaaccttcaagacct | *Alfa-gt* | 4.7×10^-7^ | T4 (hmC) |
| pLbCas12a-crRNA (*gp56*) | atcaccgagaatatcccagtatt & gagctggtcttcgggggacattt | *gp56* | 2.0×10^-6^ | T4 (C) |
| pLbCas12a-crRNA (*IPI*) | taccattaccgaagctactctta & gcaataatcttgagatgtgccgc | *IPI* | 3.4×10^-5^ | T4 *ΔIPI* |

**Table S7.** Primers used to amplify defense systems

| **#** | **Sequence (5' to 3')** | |
| --- | --- | --- |
| hhe | Fwd | cctgcaggtctcgggctattcgcGTGGCAACTGCTGGGATAATG |
|  | Rev | agcgatccgtcccgagatagggtCTTGCTATGATTCGGCTGATG |
| Druantia type III | Fwd | cctgcaggtctcgggctattcgcCATTCGGCATCACGAATAACC |
|  | Rev | agcgatccgtcccgagatagggtCCTACAATGCTTGTAATGAATGCTG |
| Gabija | Fwd | ctgcaggtctcgggctattcCTATGTCTGCATGAACATCGTTAC |
|  | Rev | gatccgtcccgagatagggtGCTTTGCGAATATTGGATTCAATG |
| Septu | Fwd | ctgcaggtctcgggctattcGCAGTTCTTCGATGTTCATCATC |
|  | Rev | gatccgtcccgagatagggtGATATGATGATTTCGAACATTCCAG |
| Zorya type II | Fwd | ctgcaggtctcgggctattcCCATTGAAGATGTTCTGGTTAAC |
|  | Rev | gatccgtcccgagatagggtACTCTGTATCAGTACAACGGTAAG |
| SspBCDE | Fwd | ctgcaggtctcgggctattcgcCACGAGAGTCAGTATTAAGGCA |
|  | Rev | agcgatccgtcccgagatagggtACGATGATCTCGCTCCTATCA |
| AVAST type4 | Fwd | ctgcaggtctcgggctattcgcCTAACACTTCCACATAGAAATTCC |
|  | Rev | agcgatccgtcccgagatagggtCTAAGCATCTGGACAATAGAGAC |
| Shedu | Fwd | ctgcaggtctcgggctattcgcCTTCAAGGAAGCTCTGAGGC |
|  | Rev | agcgatccgtcccgagatagggtGAAGTCGCTGTCGTTCTCAA |
| Retron Ec78 | Fwd | ctgcaggtctcgggctattcgcGTTTCATGATACACGCTAGACC |
|  | Rev | agcgatccgtcccgagatagggtAACCACATCTTAATTCAATGTCG |
| qatABCD | Fwd | ctgcaggtctcgggctattcgcACAAAGAACATTTGACAGCTC |
|  | Rev | agcgatccgtcccgagatagggtAAGAGCTGAACCTGACACTG |
| ppl | Fwd | ctgcaggtctcgggctattcgcTGATGCGTGCACTACGCAAA |
|  | Rev | agcgatccgtcccgagatagggtTGACGAATAAACAACAGCTGGC |
| mzaABCDE | Fwd | ctgcaggtctcgggctattcgcTTGGTCAGAATATCGTCCAGC |
|  | Rev | agcgatccgtcccgagatagggtGGTACGTCTGATCGATAGGGA |
| Restriction-like | Fwd | ctgcaggtctcgggctattcgcCAGATGAGCAGTCATATTGTAGTGG |
|  | Rev | agcgatccgtcccgagatagggtAGATCACCCTGCGATTGCAAAG |
| Retron Ec67 | Fwd | ctgcaggtctcgggctattcgcGGAAGTCGCTTAACGTATTG |
|  | Rev | agcgatccgtcccgagatagggtCATACTTAGATGTCCGTCTGT |

**Table S8.** Sequence of the pSEC1 vector backbone

**Sequence (5' to 3')**

accctatctcgggacggatcgcttcatgtggcaggagaaaaaaggctgcaccggtgcgtcagcagaatatgtgatacaggatatattccgcttcctcgctcactgactcgctacgctcggtcgttcgactgcggcgagcggaaatggcttacgaacggggcggagatttcctggaagatgccaggaagatacttaacagggaagtgagagggccgcggcaaagccgtttttccataggctccgcccccctgacaagcatcacgaaatctgacgctcaaatcagtggtggcgaaacccgacaggactataaagataccaggcgtttccccctggcggctccctcgtgcgctctcctgttcctgcctttcggtttaccggtgtcattccgctgttatggccgcgtttgtctcattccacgcctgacactcagttccgggtaggcagttcgctccaagctggactgtatgcacgaaccccccgttcagtccgaccgctgcgccttatccggtaactatcgtcttgagtccaacccggaaagacatgcaaaagcaccactggcagcagccactggtaattgatttagaggagttagtcttgaagtcatgcgccggttaaggctaaactgaaaggacaagttttggtgactgcgctcctccaagccagttacctcggttcaaagagttggtagctcagagaaccttcgaaaaaccgccctgcaaggcggttttttcgttttcagagcaagagattacgcgcagaccaaaacgatctcaagaagatcatcttattaaggggtctgacgctcagtggaacgaaaactcacgttaagggattttggtcatgagattatcaaaaaggatcttcacctagatccttttaaattaaaaatgaagttttaaatcaatctaaagtatatatgagtaaacttggtctgacagttaccaatgcttaatcagtgaggcacctatctcagcgatctgtctatttcgttcatccatagttgcctgactccccgtcgtgtagataactacgatacgggagggcttaccatctggccccagtgctgcaatgataccgcgagacccacgctcaccggctccagatttatcagcaataaaccagccagccgattcgagctcgccccggggatcgaccagttggtgattttgaacttttgctttgccacggaacggtctgcgttgtcgggaagatgcgtgatctgatccttcaactcagcaaaagttcgatttattcaacaaagccgccgtcccgtcaagtcagcgtaatgctctgccagtgttacaaccaattaaccaattctgattagaaaaactcatcgagcatcaaatgaaactgcaatttattcatatcaggattatcaataccatatttttgaaaaagccgtttctgtaatgaaggagaaaactcaccgaggcagttccataggatggcaagatcctggtatcggtctgcgattccgactcgtccaacatcaatacaacctattaatttcccctcgtcaaaaataaggttatcaagtgagaaatcaccatgagtgacgactgaatccggtgagaatggcaaaagcttatgcatttctttccagacttgttcaacaggccagccattacgctcgtcatcaaaatcactcgcatcaaccaaaccgttattcattcgtgattgcgcctgagcgagacgaaatacgcgatcgctgttaaaaggacaattacaaacaggaatcgaatgcaaccggcgcaggaacactgccagcgcatcaacaatattttcacctgaatcaggatattcttctaatacctggaatgctgttttcccggggatcgcagtggtgagtaaccatgcatcatcaggagtacggataaaatgcttgatggtcggaagaggcataaattccgtcagccagtttagtctgaccatctcatctgtaacatcattggcaacgctacctttgccatgtttcagaaacaactctggcgcatcgggcttcccatacaatcgatagattgtcgcacctgattgcccgacattatcgcgagcccatttatacccatataaatcagcatccatgttggaatttaatcgcggcctcgagcaagacgtttcccgttgaatatggctcataacaccccttgtattactgtttatgtaagcagacagttttattgttcatgatgatatatttttatcttgtgcaatgtaacatcagagattttgagacacaacgtggctttccccccccccccctgcaggtctcgggctattcgc
